# Glutamine addiction is a therapeutic target to block emergency myelopoiesis

**DOI:** 10.64898/2026.03.26.714544

**Authors:** Oakley C. Olson, Ruiyuan Zhang, Melissa A. Proven, James W. Swann, Kexuan Huang, William E. Lowry, Emmanuelle Passegué

## Abstract

Inflammation-driven emergency myelopoiesis (EM) contributes to the progression of many solid cancers and inflammatory diseases, yet therapeutic strategies to selectively suppress EM without compromising hematopoiesis remain lacking. Here, we use functional and single-cell transcriptomic analyses to determine metabolic programs organizing the hematopoietic hierarchy, myeloid lineage commitment, and myeloid differentiation. We identify de novo glutamine biosynthesis as a stem cell-specific survival mechanism allowing independence from exogenous glutamine. We show that myeloid differentiation is characterized by Myc-driven upregulation of mitochondrial respiration, which is hyperactivated during EM and renders myeloid progenitors dependent on glutaminolysis to fuel the TCA cycle. Both genetic and pharmacologic targeting of glutaminase suppresses EM and impairs breast tumor progression by reducing intratumoral neutrophil infiltration. Our study defines a central role for Myc-glutaminolysis in driving EM, identifies glutaminolysis as a therapeutic target to normalize maladaptive EM, and highlights myeloid overproduction as a systemic problem requiring HSPC targeting.

**HIGHLIGHTS:**

- HSC survival depends on de novo glutamine biosynthesis via glutamine synthetase
- Myc hyperactivation drives mitochondrial biogenesis during emergency myelopoiesis
- Myeloid progenitors become glutamine-addicted to fuel Myc-driven TCA cycle activity
- Glutaminase deficiency in HSPCs blunts tumor-promoting neutrophil production

**ETOC BLURB:** Olson et al. show that emergency myelopoiesis, the inflammatory overproduction of myeloid cells that drives regeneration, depends on Myc-driven mitochondrial respiration and glutamine addiction in hematopoietic progenitors. Targeting glutaminase in hematopoietic stem and progenitor cells suppresses pathological myelopoiesis, reduces tumor-promoting neutrophil production, and slows breast tumor growth.

## INTRODUCTION

Continuous production of myeloid cells by self-renewing HSCs located in the bone marrow (BM) is critical to maintaining organismal health^1–3^. The hematopoietic system must be able to meet dynamic changes in myeloid cell demands, such as in response to infection or chemotherapeutic treatment. However, overproduction of myeloid cells is implicated in the pathogenesis of many solid cancers and inflammatory diseases^1^. Indeed, tumor-associated myeloid cells are particularly potent drivers of cancer progression, but clinical interventions that target them have been largely unsuccessful^4^. Blocking emergency myelopoiesis (EM) pathways, the mechanisms by which inflammatory signaling redirects the hematopoietic system toward an amplified myeloid output, has been proposed as an attractive therapeutic strategy to normalize myelopoiesis without compromising innate immune system function^5,6^. However, despite our developing understanding of EM regulation, we lack effective strategies to target these pathways.

HSCs produce mature myeloid cells through sequential epigenetic and transcriptional alterations that drive expansion, lineage commitment, and terminal differentiation along a committed myeloid trajectory in the BM, involving myeloid-biased multipotent progenitor 3 (MPP3) and granulocyte/macrophage progenitors (GMP), which is amplified during EM^7^. Metabolic adaptation is now recognized as a major driver of functional plasticity in both normal HSCs and HSC-derived leukemic stem cells (LSC)^2,3^. HSC quiescence depends on glycolysis and fatty acid (FA) oxidation^8,9^, with protective autophagy preventing activation-induced mitochondria accumulation to maintain HSC lifelong function^10,11^. During aging, inflammation drives glycolytic impairment, which fasting/refeeding can reset to restore HSC regenerative potential^12^. Mitochondrial respiration defects impair HSC differentiation but not their proliferation and self-renewal, while blocking mitochondrial pyruvate metabolism selectively impacts steady state myelopoiesis but not EM^13^. Metabolic fuel flexibility is also observed as a dynamic adaptation in malignant myeloid leukemias and contributes to their pathogenic behavior^2^. Both upregulation of FA oxidation and nicotinamide metabolism have emerged as critical resistance mechanisms to Venetoclax/Azacitidine treatment, which preserve LSCs^14,15^, while glutamine metabolism has been found critical for maintaining minimal residual disease (MRD) following the stress of chemotherapeutic treatment, with inhibitors of glutamine metabolism being investigated as anti-leukemic agents^16^. Despite our developing understanding of metabolic regulation of HSC and LSC function, it remains poorly understood how the metabolism of the hematopoietic system adapts to produce myeloid cells under steady-state and stress conditions, particularly in myeloid-biased progenitors. Here, we leveraged functional and single-cell transcriptomic approaches to identify metabolic programs that organize the hematopoietic hierarchy, drive myelopoiesis, and control EM in normal and maladaptive conditions.

## RESULTS

### Mapping HSPC metabolic programs

To understand the regulation of metabolism during myeloid lineage commitment and differentiation, we mapped the HSPC compartment by single-cell RNA sequencing (scRNA-seq) using a combination of Lin^-^/c-Kit^+^ (LK) and Lin^-^/Sca-1^+^/c-Kit^+^ (LSK) BM populations (**Figure 1A**; **Figure S1A**). Uniform Manifold Approximation and Projection (UMAP) dimensional reduction and HemaScribe cell annotation^17^ resolved all major hematopoietic lineages and identified HSPC populations (**Figure 1B**). Strikingly, UMAP dimensional reduction using only the metabolic genes (1,647 out of 22,021 genes, ∼ 7%) demonstrated that unique metabolic transcriptional programs alone can distinguish the hematopoietic lineages in transcriptional space (**Figure 1C**). Interestingly, of the 1,647 metabolic genes, 1,514 (∼ 92%) were identified as highly variable genes in our dataset, illustrating their importance in the global data structure. Differentiation from HSCs into myeloid and erythroid lineages was broadly associated with increased oxidative phosphorylation (OXPHOS), pentose phosphate pathway (PPP), and glutathione metabolism, with increased glutathione recycling specific to myeloid differentiation based on geneset enrichment analysis (GSEA) and pathway-specific gene expression analyses (**Figure S2A and S2B**). To dissect metabolic pathway usage along lineage trajectories, we employed a graph neural network model to estimate cell-wise metabolic flux^18^, observing increased estimated flux through the TCA cycle, polyamine and pyrimidine metabolisms, and amino acid (AA) uptake in myeloid and erythroid/megakaryocyte progenitors, while increased glycolysis and PPP activity was specific to myeloid progenitors (**Figure 1D**). Consistent with this, metabolomic analysis of purified LSK and GMP populations showed a broad enrichment in metabolites associated with a biosynthetic and metabolically active cell state in GMPs, including glutamine, nucleotides and TCA intermediates (**Figure S2C**). Measurement of mitochondrial membrane potential with TMRE dye and reduced glutathione with ThiolTracker confirmed the increase in oxidative metabolism and redox buffering capacity in MPP3s and GMPs relative to HSCs (**Figure S2D**). Collectively, these data illustrate the metabolic activation of committed progenitors downstream of HSCs and specific expansion of redox buffering capacity in myeloid progenitors through increased PPP-dependent regenerative glutathione metabolism.

**Figure 1.**
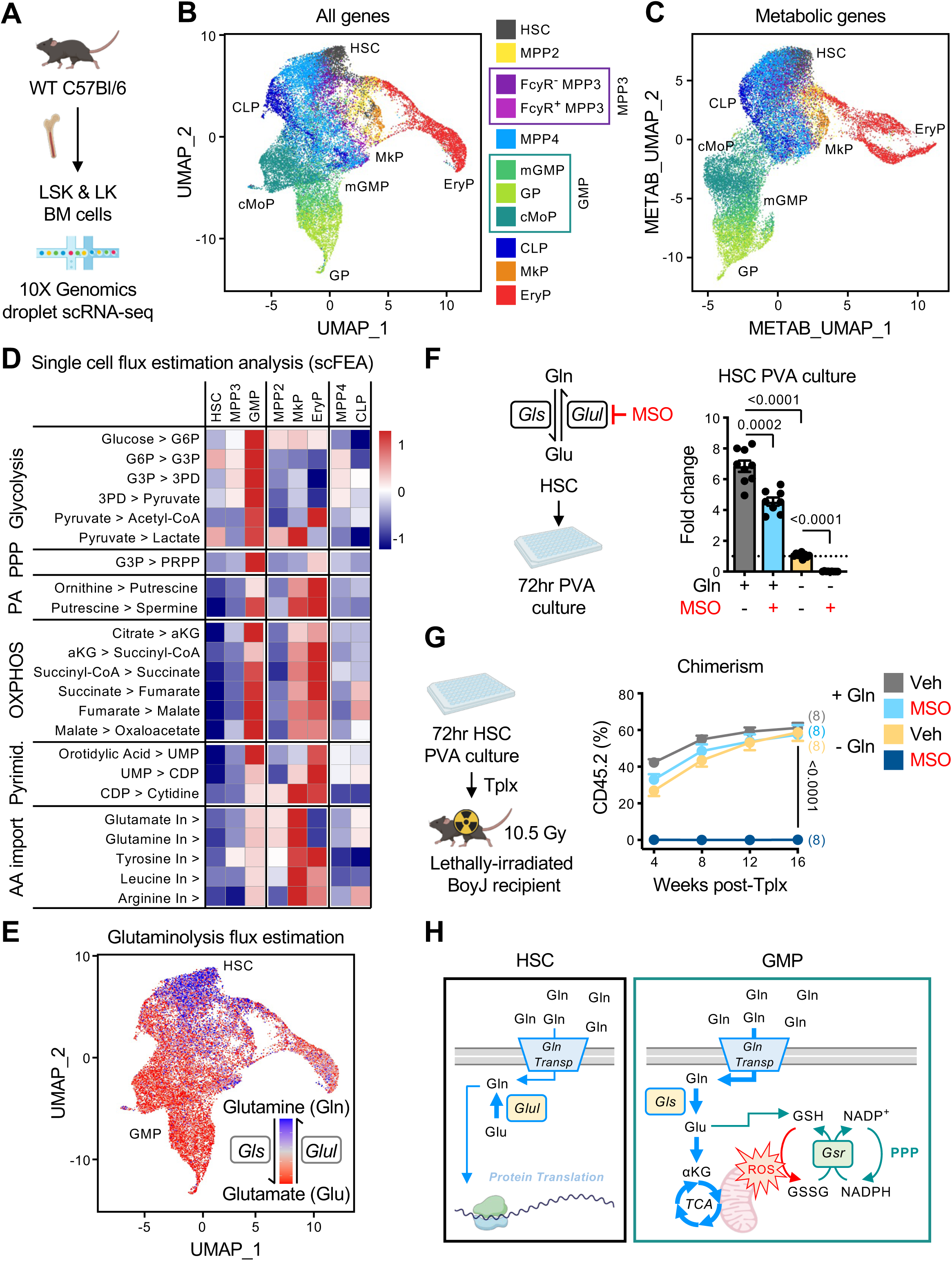
Hematopoiesis is organized by metabolic transcriptional programs. (A) Transcriptional regulation of steady-state HSPCs analyzed by scRNA-seq of BM LSK and LK populations. (B) UMAP representation of HSPCs using HemaScribe cell type identification. (C) UMAP representation of HSPCs using only metabolic genes. (D) Single-Cell Flux Estimation Analysis (scFEA) of indicated metabolic pathways activity in distinct HSPC populations. PPP, pentose phosphate pathway; PA, polyamine metabolism; Pyrimid, pyrimidine nucleotide biosynthesis; AA import, amino acid import. (E) Feature plot of the metabolic flux module of glutamine to glutamate conversion. (F-G) HSC response to glutamine starvation following 72 hours PVA culture with or without glutamine (±Gln) and the glutamine biosynthesis inhibitor methionine sulfoximine (±MSO): (F) expansion rates in culture expressed as fold change of plated cells (n=8-9; 3 independent experiments); and (G) engraftment rates following transplantation (TPLX) in lethally irradiated congenic recipient mice (n=7-8; 2 independent experiments). (H) Scheme representing the dependency of HSCs to de novo glutamine biosynthesis to maintain protein translation rates, and of myeloid lineage GMPs to glutaminolysis to supports increased OXPHOS and pentose phosphate pathway to maintain redox balance. Data are means ± S.E.M. (F, G); circles represent individual mice (F); *P. values* were obtained by an unpaired t-test (F, G). See also Figures S1 and S2.

### HSC survival depends on preserving the intracellular glutamine pool

While expression of metabolic transcriptional programs was broadly suppressed in HSCs, we observed a specific enrichment of glutamine biosynthesis relative to glutaminolysis by single-cell flux estimation (**Figure 1E**). This was the result of enriched expression of glutamine synthetase (*Glul*), which catalyzes glutamine biosynthesis from glutamate and ammonia, in HSCs compared to MPP3s and GMPs (**Figure S2E and S2F**). Glutamine is the most abundant amino acid, and most cells depend on the catabolism of exogenous glutamine for their proliferation and survival^19^. We therefore sought to determine whether *Glul* expression in HSCs could render them independent of exogenous glutamine, similar to other cells known to express *Glul*, such as embryonic stem cells (ESC)^20^. HSCs cultured *ex vivo* in self-renewal conditions expanded ∼ 6-fold over 72 hours, while glutamine starvation produced a cytostatic effect without cell expansion but with maintenance of long-term engraftment upon transplantation into lethally irradiated recipient mice (**Figure 1F and 1G**). This was specific to HSCs as glutamine starvation in MPP3 was cytotoxic (**Figure S2G**), and dependent on Glul activity because inclusion of the specific inhibitor methionine sulfoximine (MSO)^20^ caused HSC death (**Figure 1F**). Indeed, MSO treatment eliminated phenotypic HSCs (Lin^-^/Sca-1^+^/c-Kit^+^/CD48^-^/CD150^+^) during glutamine starvation and completely abrogated long-term engraftment potential (**Figure 1G**; **Figure S1B and S2H**). Notably, measurement of protein translation rate by OP-Puro incorporation showed that Glul activity was essential to maintain proteostasis during glutamine starvation, with a major decrease in OP-Puro incorporation in MSO-treated HSCs similar to mTOR inhibition by rapamycin (**Figure S2J**). These results establish that HSCs prioritize utilization of glutamine for protein translation rather than as a fuel source to meet biosynthetic and bioenergetic demand, and the ability to biosynthesize glutamine as an important cytoprotective mechanism specific to HSCs that is required for their survival under stress (**Figure 1H**).

### Myc hyperactivation is a driver of EM

Next, we sought to understand the metabolic adaptation of the hematopoietic system following EM engagement. We treated mice with a single dose of 5-fluorouracil (5FU) as a prototypic model of regenerative EM, in which myeloablative chemotherapy clears the BM and allows tracing of myeloid regeneration along a typified kinetic involving HSC activation, MPP3 amplification, and GMP cluster formation^21^. We profiled BM HSPCs by scRNA-seq analysis at day 0 (D0) steady-state (using LK + LSK cells) and during the early phase of the regenerative process at day 7 to day 10 (D7-10) post-5FU treatment (using LK cells) (**Figure 2A and 2B**). Single cell flux estimation indicated increased glycolysis, TCA cycle, polyamine and pyrimidine metabolisms, as well as increased AA uptake in all HSPCs on D7 and D8 (**Figure S3A**). Searching for drivers of this metabolic adaptation, GSEA of D8 GMPs identified upregulation of Myc transcriptional programs as a dominant feature of EM activation (**Figure 2C**), with an acute upregulation of Myc transcriptional response throughout the HSC/MPP3/GMP hierarchy during EM (**Figure 2C**). Accordingly, using a knock-in *Myc-eGFP* reporter mouse^22^, we observed clear upregulation of Myc protein in D8 MPP3s and GMPs (**Figure S3B**). To determine whether Myc hyperactivation was necessary for EM, we crossed the HSC-specific *Scl-Cre^ER^* conditional Cre diver mice^23^ with Myc floxed mice^24^ to excise a single copy of *Myc* in *Scl-Cre^ER^:Myc^+/flox^* (*Myc^+/Δ^*) mice upon tamoxifen injection (**Figure 2D**; **Figure S3C**). Hematopoietic-specific *Myc* haploinsufficiency had no impact on BM cellularity or neutrophil counts one month after recombination compared to tamoxifen-treated *Myc^+/flox^* control (Ctrl) mice (**Figure S3D**). We also confirmed that deletion of both *Myc* alleles resulted in aplastic anemia and quick death of the animals, as previously reported^25^. Upon challenge with 5FU one month after recombination, *Myc^+/Δ^* mice recovered circulating blood counts of neutrophils with similar kinetics as Ctrl mice but without the neutrophil overproduction characteristic of EM engagement^21^ (**Figure 2E**). This was specific to the myeloid lineage since erythroid regeneration was unaffected in 5FU-treated *Myc^+/Δ^* mice (**Figure 2E**). In the BM of 5FU-treated *Myc^+/Δ^* mice, we found prolonged expansion of HSCs at D12 (**Figure S3E**), which was suggestive of impaired differentiation and increased self-renewal expansion, and reduced levels of late myeloid progenitors (granulocyte progenitors or GPs, a neutrophilic-committed subset of GMPs) at D14 (**Figure 2F**) that correlated with hypotrophic GMP patches at D10 and smaller GMP clusters at D12 (**Figure 2G**). In addition, the expansion of immature CD101- neutrophils^26^ at D14, characteristic of EM engagement, was not observed in the BM of 5FU-treated *Myc^+/Δ^* mice (**Figure S1A and S3F**). Collectively, these data establish the essential role of Myc hyperactivation for a productive EM response, including robust GMP cluster formation and enhanced output of immature neutrophils, which is compromised upon Myc haploinsufficiency

**Figure 2.**
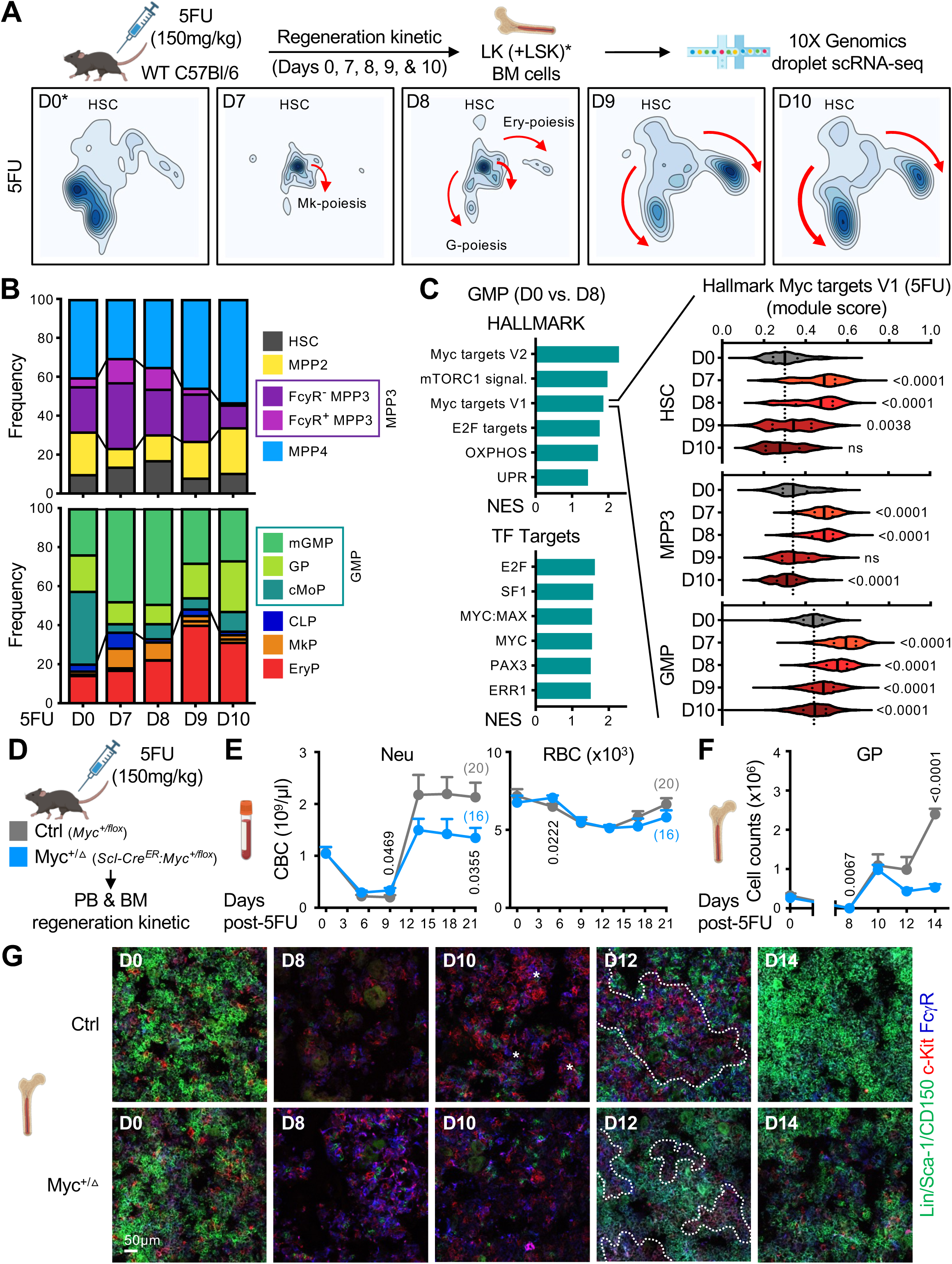
Emergency myelopoiesis is driven by Myc hyperactivation. (A) Transcriptional regulation of regenerative HSPCs following 5FU myeloablation analyzed by scRNA-seq of BM LSK and LK populations (top), with contour density plots showing successive reconstitution of hematopoietic lineages during regeneration (bottom). (B) Quantification of HSPCs and lineage-committed progenitors upon HemaScribe identification. (C) Hallmark and TFT legacy GSEA analysis of significant (*p* <0.05) transcriptional changes between GMP at day 0 (D0) steady state and day 8 (D8) post-5FU treatment (left), and single-cell assessment of Hallmark Myc V1 transcriptional signature in HSC, MPP3, and GMP populations during the regeneration kinetic (right). NES, normalized enrichment score. (D-G) Regenerative response in hematopoietic-specific *Myc* haplo-insufficient mice: (D) scheme of 5FU treatment in control (Ctrl) and hematopoietic-specific conditionally deleted Myc mice (*Myc^+/Δ^*); quantification of (E) peripheral blood neutrophils (Neu) and RBCs (n=16-20; 2 independent experiments), and (F) BM granulocyte progenitors (GP) (n=3-7 mice per time point and genotype; 3 independent experiments) post-5FU treatment; and (G) immunofluorescence imaging of BM GMP patches (stars) and GMP clusters (dotted line) post-5FU treatment (representative images from 3 independent experiments). Data are violin plots with quartiles (C, right) or means ± S.E.M. (E, F); *P. values* were obtained by the Kolmogorov-Smirnov test (C, left), Kruskal-Wallis test (C, right), or unpaired t-test (E, F). See also Figures S3.

### Myc-driven mitochondrial respiration during EM

To understand how Myc drives EM activation, we profiled BM HSPCs by scRNA-seq analysis of LK/LSK cells isolated from Ctrl and *Myc^+/Δ^* mice at D0 and D8 post-5FU treatment (**Figure 3A**; **Figure S4A**). Consistent with its prominent overexpression after 5FU treatment, Myc haploinsufficiency led to global suppression of EM transcriptional changes (**Figure S4B**), with suppression in D8 HSCs of coordinated metabolic programs necessary for activation and proliferation, including OXPHOS, nucleotide metabolism and ribosome biogenesis (**Figure S4C**). In D8 MPP3, we observed the same Myc-dependent alterations as in HSC, with additional downregulated biosynthetic metabolic programs including FA biosynthesis, AA metabolism, and spliceosome regulation of mRNA (**Figure S4C**). Single cell flux estimation demonstrated that Myc haploinsufficiency blunted the metabolic adaptation observed in response to 5FU across the HSC/MPP3/GMP hierarchy, with a more prominent effect on MPP3s (**Figure 3B**). Specifically, the upregulation of a Myc-regulated mitochondrial biogenesis transcriptional program^27^ observed in Ctrl HSPC populations at D8 was significantly reduced in all *Myc^+/Δ^* HSPC populations (**Figure 3C**). To directly assess the impact of this change, we isolated HSCs, MPP3s, and LSK BM cells at D8 and GMPs at D10 post-5FU treatment to measure mitochondrial content by Tomm20 immunofluorescence and OXPHOS by Seahorse analyses. While 5FU-driven mitochondrial biogenesis was only blunted in D8 *Myc^+/Δ^* HSCs (**Figure S4D**), it was completely blocked in D8 MPP3s and D10 GMPs, with no significant difference between steady state and 5FU treatment in *Myc^+/Δ^* mice (**Figure 3D and 3E**). Moreover, measurement of mitochondrial membrane potential showed no increase in oxidative metabolism in D8 *Myc^+/Δ^* HSCs (**Figure S4E**), and extracellular flux analysis indicated a loss in D8 *Myc^+/Δ^* LSK cells of the increased maximal respiration characteristic of D8 Ctrl LSK cells (**Figure 3F**), both confirming Myc requirement in HSCs for increased OXPHOS upon EM activation. To further understand the regulation of Myc hyperactivation during EM, we performed single-cell Multiome analyses combining both ATAC and RNA-seq measurements on D0 (LK + LSK cells) and D8 (LK cells) HSPCs from Ctrl mice (**Figure S5A and S5B**). We found significant alterations in the chromatin landscape in response to EM activation, with massive opening of chromatin in response to 5FU and increased accessibility of AP-1 factors in MPP3s and CCAAT/enhancer-binding (C/EBP) proteins in GMPs (**Figure S5C to S5E**). Despite the upregulation of Myc transcriptional targets, we did not observe increased chromatin accessibility of Myc motifs in any HSPC population (**Figure S5D**). Rather, Myc motifs appear readily accessible at baseline (**Figure S5E**), indicating that HSPCs are primed for Myc hyperactivation without requiring changes in chromatin accessibility. Taken together, these results demonstrate that Myc is a driver of metabolic adaptation in EM, with the HSPC hierarchy poised to quickly stimulate mitochondrial biogenesis and OXPHOS to meet the energetic demands of enhanced myeloid cell production.

**Figure 3.**
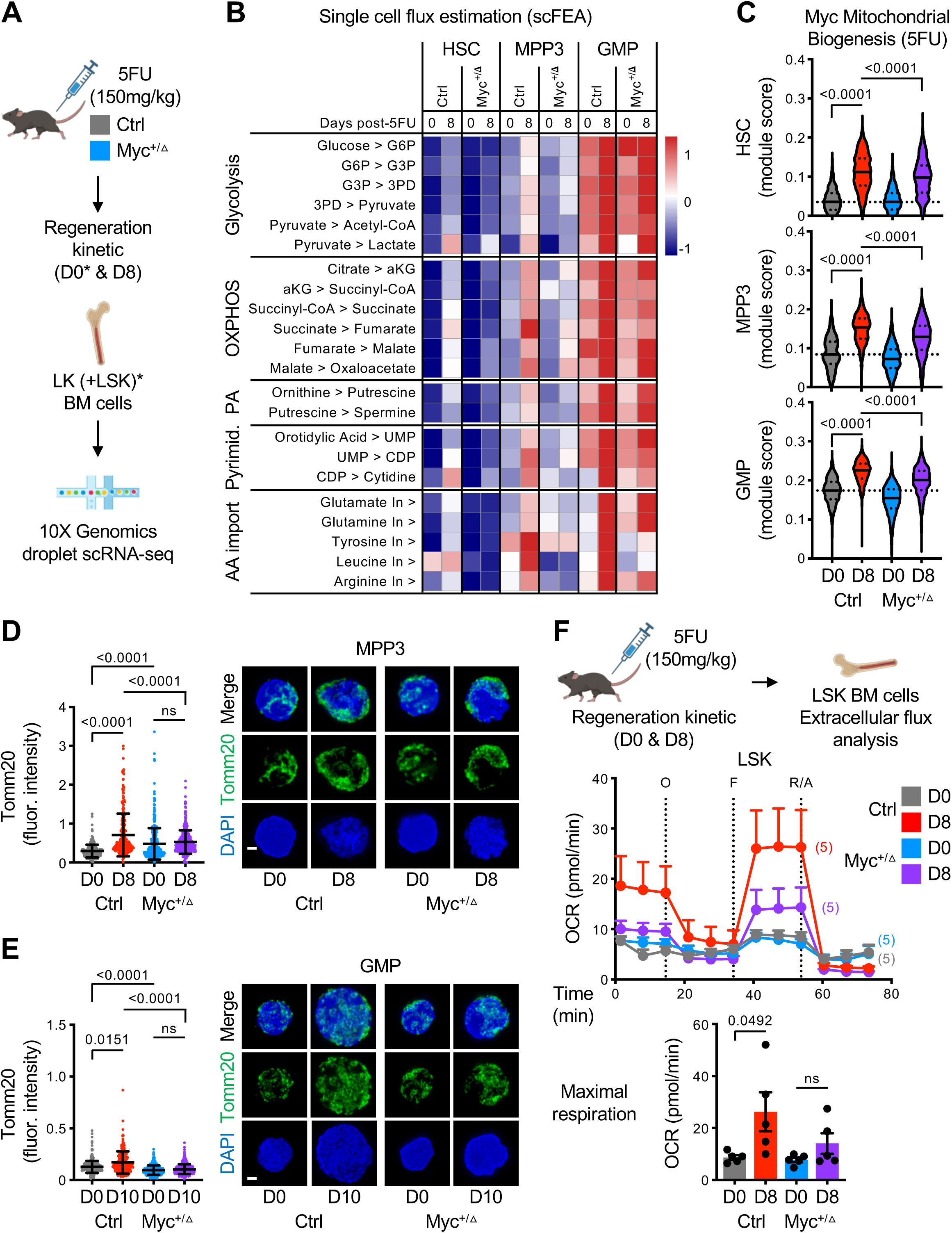
Myc-dependent increase in mitochondrial biogenesis and oxidative phosphorylation sustains emergency myelopoiesis. (A-C) Transcriptional regulation of regenerative Ctrl and *Myc^+/Δ^* HSPCs at day 0 (D0) and day 8 (D8) post-5FU analyzed by scRNA-seq of BM LSK and LK populations: (A) scheme; (B) scFEA analysis of indicated metabolic pathways; and (C) single-cell assessment of Myc mitochondrial biogenesis score^27^ in HSC, MPP3, and GMP populations. PA, polyamine metabolism; Pyrimid, pyrimidine nucleotide biosynthesis; AA import, amino acid import. (D-E) Mitochondria quantification in regenerative Ctrl and *Myc^+/Δ^* (D) MPP3 (n=269-499 cells; 3 independent experiments) and (E) GMP (n=195-919 cells; 3 independent experiments) at D0 and D8 post-5FU analyzed by Tomm20 immunofluorescence, with quantification of Tomm20 fluorescence (fluor.) intensity (left) and representative image (selected from 3 independent experiments; right). (F) Scheme of OXPHOS measurement by extracellular flux analysis in regenerative Ctrl and *Myc^+/Δ^* LSK at D0 and D8 post-5FU (top), with oxygen consumption rates (OCR) levels (middle) and detailed maximal respiration levels (bottom) (n=5; 4 independent experiments). O, Oligomycin A; F, FCCP; R/A, Rotenone/Antimycin A. Data are violin plots with quartiles (C) or means ± S.D. (D, E) or ± S.E.M. (F); dots represent individual cells (D, E) and circles individual mice (F); *P. values* were obtained by the Kruskal-Wallis test (C) or an unpaired t-test (D, E, F). See also Figures S4 and S5.

### Glutaminolysis drives EM

To identify potential metabolic dependencies underlying this adaptation, we next sought to determine the specific TCA fuel requirements for EM. We cultured isolated Ctrl LSK and GMP cells under EM-like myeloid expansion conditions with inhibitors blocking the utilization of glutamine, FA, or pyruvate by the TCA cycle, and inhibition of the electron transport chain by Rotenone^28^ as a positive control (**Figure 4A**). Strikingly, treatment with the glutaminolysis inhibitor BPTES^29^ led to a ∼4-fold reduction in LSK and ∼2-fold reduction in GMP cell expansion (**Figure 4B**). In contrast, no effect was observed with inhibition of FA oxidation with Etomoxir^30^ or pyruvate uptake with UK5099^31^, demonstrating that utilization of glutamine is required specifically during myeloid expansion. In fact, similar cultures performed with *Myc^+/Δ^* cells showed that Myc-driven myeloid expansion was largely dependent on glutaminolysis for LSK expansion, and entirely dependent for GMP expansion (**Figure S6A**). To assess anaplerotic glutamine metabolism in HSPCs, we then performed 1-hour stable isotope tracing of ^13^C_5_-glutamine in Ctrl mice at D0 and D8 post-5FU treatment (**Figure S6B**). As expected, we found glutamine-derived ^13^C labelling in TCA cycle metabolites and amino acids in isolated LK cells (**Figure S6B and S6C**). To determine the functional role of EM glutaminolysis *in vivo*, we employed a genetic approach and crossed the HSC specific *Scl-Cre^ER^*driver mice with glutaminase (*Gls*) floxed mice^32^ to remove both copies of the rate-limiting enzyme of glutaminolysis (**Figure 4C**). Hematopoietic-specific loss of glutaminase in tamoxifen-treated *Scl-Cre^ER^:Gls^flox/flox^* (*Gls^Δ/Δ^*) mice had no impact on BM neutrophils 1 month after recombination compared to tamoxifen-treated *Gls^flox/flox^* Ctrl mice (**Figure 4D**). In contrast, upon 5FU treatment, we observed delayed recovery of circulating blood neutrophils, without the overproduction characteristic of EM induction (**Figure 4E**). Again, this was specific to myeloid regeneration without impairment of erythropoiesis (**Figure 4E**). In the BM, 5FU-treated *Gls^Δ/Δ^* mice also showed delayed recovery of late myeloid progenitors (GP) and delayed formation and differentiation of GMP clusters, ultimately leading to impaired expansion of CD101^-^immature neutrophils (**Figure 4F and 4G; Figure S6D and S6E**). To exclude the possibility that glutaminase deficiency in neutrophil precursors rather than HSPCs was responsible for the observed phenotype, we also used the *Mrp8-Cre* driver mice^33^ to excise *Gls* in neutrophils and neutrophil progenitors in *Mrp8-Cre:Gls^flox/flox^*(*Gls^mΔ/Δ^*) mice (**Figure S6F**). However, we observed no impact on peripheral white blood cell (WBC) counts or neutrophil recovery following 5FU treatment (**Figure S6G**), demonstrating that the requirement for glutaminase during EM is restricted to HSPCs. Collectively, these results demonstrate that HSPCs rely on glutaminolysis for Myc-driven myeloid cell production during regeneration. This glutamine dependence mirrors the “glutamine-addiction” characteristic of Myc-driven cancers^19,34^, suggesting that cancers may co-opt a hyperproliferative metabolic state that is also essential for regenerative responses like EM.

**Figure 4.**
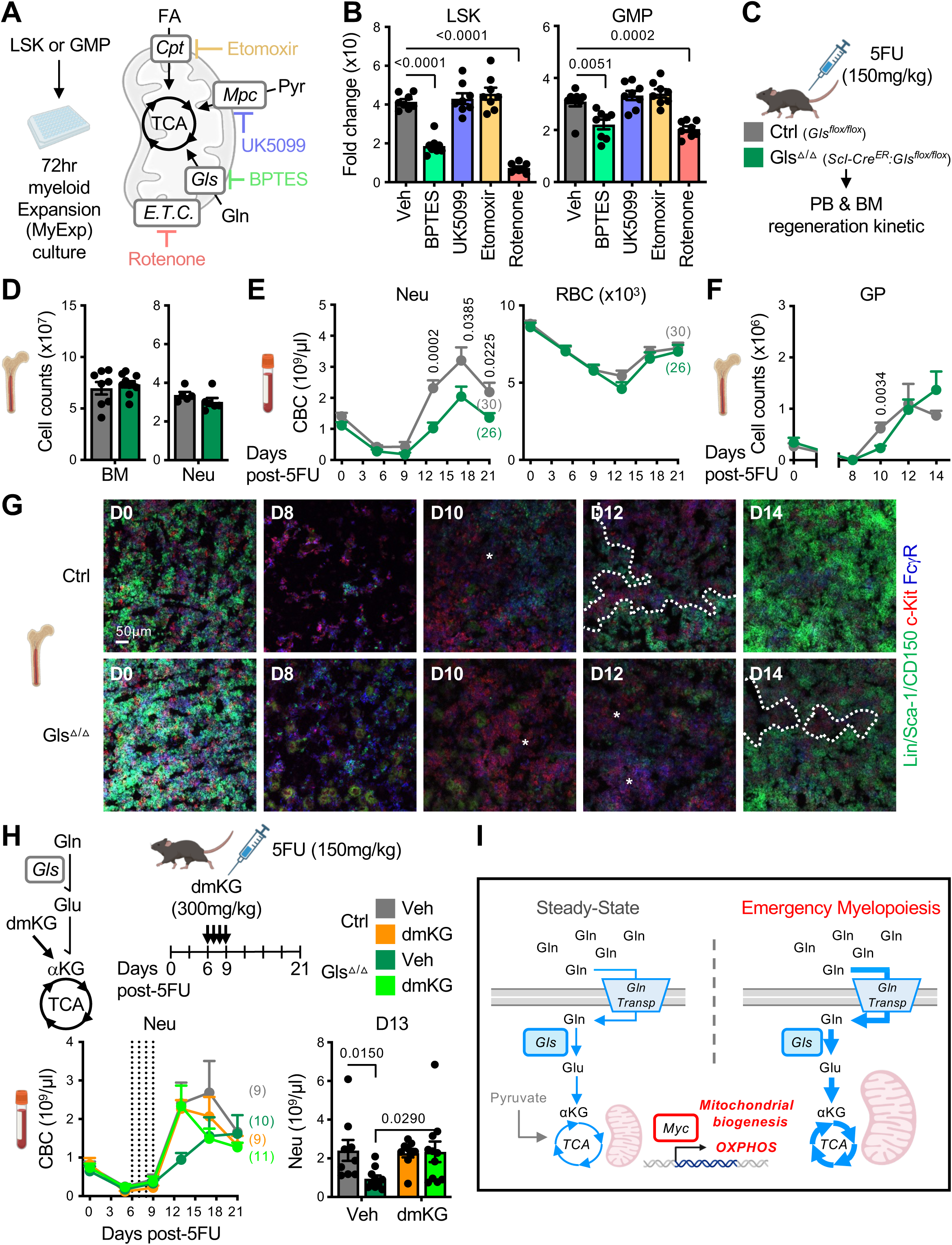
Glutamine addiction during emergency myelopoiesis. (A-B) Inhibitor-based assessment of TCA fuel dependency of HSPCs in regenerative-like myeloid expansion (MyExp) liquid culture conditions: (A) scheme of the experimental design (left) and mitochondrial targets of the inhibitors (right); and (B) quantification of LSK (left) and GMP (right) expansion after 72 hours culture (n=8; 4 independent experiments). Results are shown as fold change of plated cells. FA, fatty acid; Pyr, pyruvate; E.T.C., electron transport chain; BPT. (C-G) Regenerative response in hematopoietic-specific glutaminase deficient mice: (C) scheme of 5FU treatment in control (Ctrl) and hematopoietic-specific *Gls*-deficient mice (*Gls^Δ/Δ^*); (D) quantification of BM cellularity (n=8-9; 4 independent experiments) and neutrophils at steady state (n=4-6; 2 independent experiments); quantification of (E) peripheral blood neutrophils and RBCs (n=26-30; 3 independent experiments) and (F) BM granulocyte progenitors (GP) (n=6-16 mice per time point and genotype; 4 independent experiments) post-5FU treatment; and (G) immunofluorescence imaging of BM GMP patches (stars) and GMP clusters (dotted line) post-5FU treatment (representative images from 2 independent experiments). (H) *In vivo* rescue of impaired regeneration from glutaminase-deficient HSPCs: treatment scheme with daily dmKG injections at days 6 to 9 post-5FU treatment in Ctrl and *Gls^Δ/Δ^* mice (n=9-11; 2 independent experiments) (top), quantification of peripheral blood neutrophils (bottom), and neutrophil counts at D13 post-5FU. (I) Model of Myc-driven metabolic adaptation during emergency myelopoiesis leading to increased OXPHOS and glutamine addiction. Data are means ± S.E.M.; circles represent individual mice; *P. values* were obtained by an unpaired t-test (B, E, H). See also Figures S6 and S7.

### Anaplerotic glutamine metabolism

Having established that glutaminolysis is required for EM, we next investigated its role as an anaplerotic fuel for TCA cycle activity. We performed extracellular flux analysis on purified LSK cells from 5FU-treated Ctrl or *Gls^Δ/Δ^* mice at D8 and observed abrogation of the EM-induced increase of maximal respiration in glutaminase-deficient cells (**Figure S7A**). We next sought to determine whether supplementation with dimethyl-ketoglutarate (dmKG), a cell permeant analogue of the downstream TCA cycle metabolite α-ketoglutarate (αKG), was sufficient to rescue impaired expansion of glutaminase-deficient cells in myeloid expansion culture conditions (**Figure S7B**). For both LSK and GMP cells, supplementation with dmKG fully rescued glutaminase-deficient cell expansion. Interestingly, supplementation with the antioxidant n-acetyl cysteine (NAC) also rescued the expansion of *Gls^Δ/Δ^* GMPs, but not LSK cells (**Figure S7C**). This suggested that glutaminase is also critical to glutathione biosynthesis in GMPs, further underscoring the role of the glutathione redox system in myeloid progenitors. In addition, glutaminase deficiency did not appear to affect neutrophil immunometabolic function, either at steady state or following EM engagement, as extracellular flux analysis performed on Ctrl or *Gls^Δ/Δ^* BM neutrophils showed no changes in mitochondrial respiration or oxidative burst capacity at D0 or D14 post-5FU (**Figure S7D and S7E**). Collectively, these data show that Myc-driven activation of OXPHOS during regenerative EM leads to a specific dependence on glutamine as a critical fuel for generating TCA cycle intermediates (**Figure 4I**).

### Glutamine addiction as an EM therapeutic target

We next sought to determine whether EM glutamine addiction could be leveraged for therapeutic interventions. We previously showed in an MMTV-PyMT mouse model of breast cancer that tumor-derived G-CSF drives the activation of EM pathways and overproduction of tumor-promoting neutrophils^5^. We hypothesized that breast cancer-activated pathogenic EM could be similarly addicted to glutamine and that targeting glutaminolysis in BM HSPCs could represent a potential strategy to reduce the production of tumor-promoting neutrophils. First, we confirmed that EM induction mediated by G-CSF treatment ^21^ and triggered in the transgenic MMTV-PyMT breast cancer model both relied on Myc-driven metabolic adaptation. As expected, Myc and mitochondrial biogenesis signatures were the dominant pathways enriched in scRNA-seq analyses of HSPCs isolated from wild type (WT) mice treated daily with 5 µg of human recombinant G-CSF (**Figure S8A to S8C**). Single-cell metabolic flux estimation also identified metabolic adaptation with increased glycolysis and OXPHOS largely restricted to GMPs in G-CSF-treated mice (**Figure S8D**), consistent with recent observations regarding G-CSF effects^17^. Functionally, G-CSF-treated *Gls^Δ/Δ^* mice had suppressed myeloid cell overproduction compared to G-CSF-treated Ctrl mice (**Figure S8E and S8F**). Similarly, in the transgenic MMTV-PyMT breast cancer model, upregulation of Myc and mitochondrial biogenesis signatures were the dominant pathways enriched in scRNA-seq analyses of HSPCs isolated from tumor-bearing PyMT mice compared to WT littermate Ctrl (**Figure S9A to S9C**). Single-cell metabolic flux estimation also revealed metabolic adaptation throughout the HSPC compartment similar to regenerative EM, with particular upregulation of OXPHOS pathways (**Figure S9D**). Taken together, these results show that signatures of Myc-driven metabolic activation are consistent across EM models, particularly in GMPs.

To assess the impact of EM metabolic blockade on breast cancer progression, we orthotopically implanted a syngeneic MMTV-PyMT cell line (Py230)^35^ in *Gls^Δ/Δ^* and Ctrl mice (**Figure 5A**). Tracking tumor growth by external measurement, we observed decreased tumor volume by 8 weeks in glutaminase deficient recipient mice (**Figure 5B**). Tumor-bearing *Gls^Δ/Δ^* mice had overall decreased BM cellularity due to significantly reduced neutrophil numbers, specifically CD101^-^ immature neutrophils, and monocyte counts (**Figure 5C**; **Figure S10A**). Despite no major changes in other HSPCs, we observed a significant reduction in late myeloid progenitors (GPs) associated with the presence of hypotrophic GMP clusters in the BM of tumor-bearing *Gls^Δ/Δ^* mice (**Figure 5C and 5D; Figure S10A**). HSPC analysis by scRNA-seq confirmed that glutaminase deficiency led to a suppression of the tumor-induced GMP expansion (**Figure 5E**) and KEGG GSEA analysis of GMPs from tumor-bearing *Gls^Δ/Δ^* mice showed decreased metabolic pathways (OXPHOS, AA metabolism) and increased signs of cellular stress (p53 signaling, apoptosis) (**Figure S10B**). Extracellular flux analysis also demonstrated impaired mitochondrial respiration in both LSK cells and GMPs from tumor-bearing *Gls^Δ/Δ^* mice (**Figure 5F**; **Figure S10C**). By looking at circulating WBC and neutrophils as a function of tumor burden, we observed a reduced correlation in *Gls^Δ/Δ^* mice, demonstrating that hematopoietic glutaminase deficiency blunts tumor-induced neutrophilia during disease progression (**Figure 5G**). This inability to produce neutrophils led to a specific decrease in intratumoral neutrophils, including the tumor-educated DcTrailR1^+^ T3 neutrophils^36^ and a compensatory increase in intratumoral monocytes, but without differences in intratumoral tumor-associated macrophages (TAM) (**Figure 5H**; **Figure S1C** and **S10D**). Quantifying tumor-infiltrating leukocyte populations, we also observed a significant increase in intratumoral CD4^+^ T cells, which control breast tumor progression and metastasis^37^, but no changes in other immune populations (**Figure S10D**). Importantly, mature myeloid cell-specific glutaminase deficiency in *Gls^mΔ/Δ^* mice did not lead to any changes in tumor growth, circulating neutrophil numbers, or intratumoral pro-tumorigenic T3 neutrophils upon orthotopic Py230 implantation (**Figure 5I; Figure S10E**), demonstrating that cancer-induced EM depends on glutaminase specifically in HSPCs but not mature cells.

**Figure 5.**
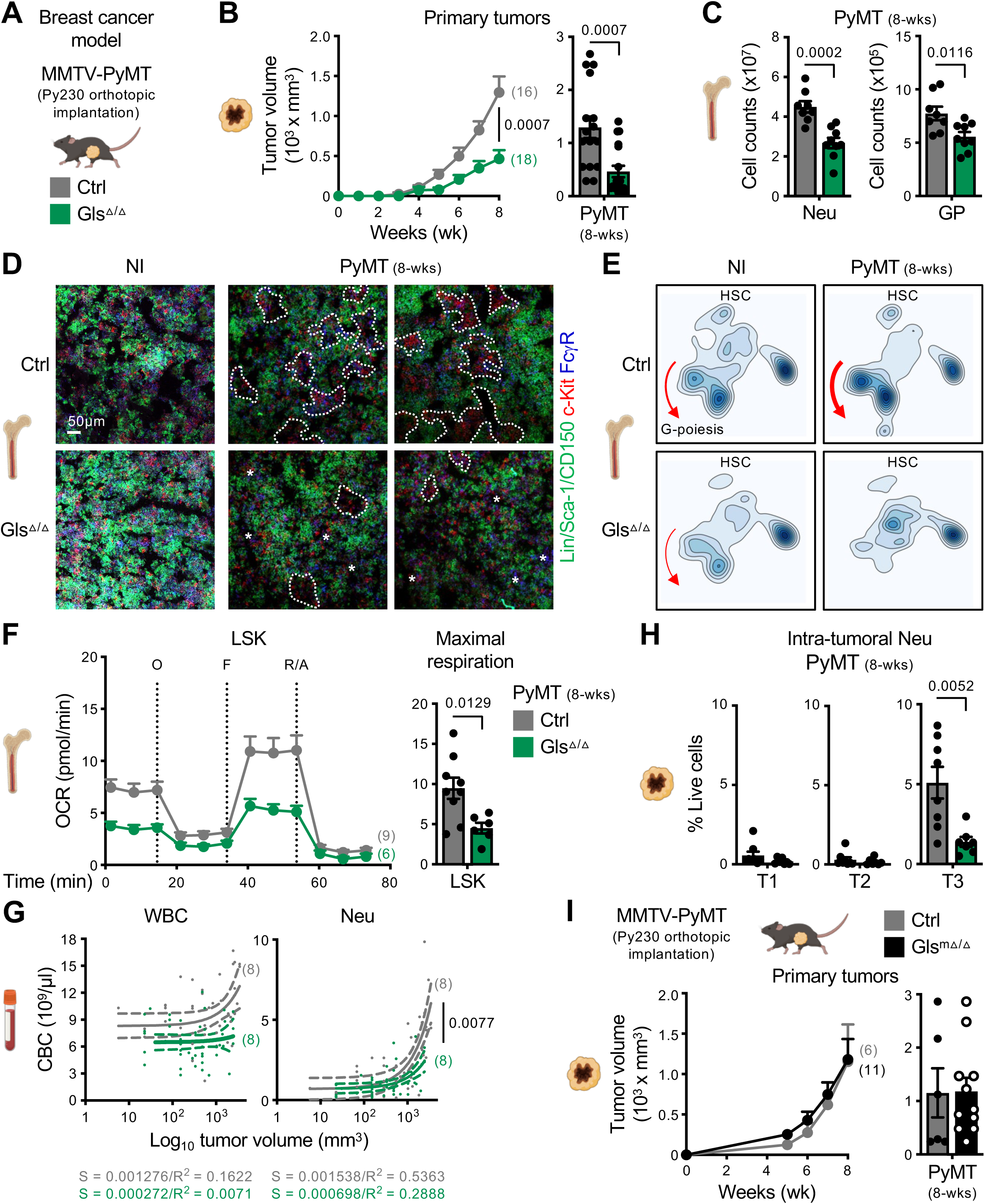
Glutamine addiction is a therapeutic target to block emergency myelopoiesis. (A-B) Targeting hematopoietic-specific glutaminolysis during breast cancer development in Ctrl and *Gls^Δ/Δ^* mice: (A) breast cancer model with orthotopic implantation of MMTV-PyMT cell line (Py230, 1×10^6^ cells); and (B) breast tumor growth over time measured by external palpation (n=16-18; 4 independent experiments; left), with tumor volume at 8 weeks (8-wks) post-orthotopic implantation (n=8-9; 2 independent experiments; right). (C-H) Characterization of non-implanted (NI) and tumor-bearing (PyMT) Ctrl and *Gls^Δ/Δ^* mice at 8-wks post-orthotopic implantation: (C) quantification of BM neutrophils (Neu) and granulocyte progenitors (GP) (n=8-9; 2 independent experiments); (D) immunofluorescence imaging of BM GMP patches (stars) and GMP clusters (dotted line) (representative images from 2 independent experiments); (E) transcriptional regulation of HSPCs analyzed by scRNA-seq, with contour density plots of BM LK cells showing tumor-induced loss of granulopoiesis in tumor-bearing *Gls^Δ/Δ^* mice; (F) OXPHOS measurement by extracellular flux analysis of LSK, with OCR levels (left) and detailed maximal respiration levels (right) (n=6-9; 6 independent experiments); (G) circulating WBC and Neu plotted as a function of PyMT tumor volume (n=16-18; 3 independent experiments), with simple linear regression reporting slope (S) and goodness of fit (R2); and (H) quantification of intratumoral pro-tumorigenic neutrophil subsets^36^ (n=8; 3 independent experiments). O, Oligomycin A; F, FCCP; R/A, Rotenone/Antimycin A; T1, DcTrailR1^-^/CD101^-^; T2, DcTrailR1^-^/CD101^+^; T3, DcTrailR1^+^/CD101^+/-^. (I) Targeting myeloid-specific glutaminolysis during breast cancer development in Ctrl and *Gls^mΔ/Δ^* mice (n=6-11; 3 independent experiments) with tumor growth over time measured by external palpation (left), and tumor volume at 8 weeks (8-wks) post-orthotopic implantation (right). Data are means ± S.E.M. (B, C, F, H, I) or linear regression with 95% C.I. (G); dots represent individual measurements (G) and circles individual mice (B, C, F, H, I); *P. values* were obtained by an unpaired t-test (B, C, F, H, I) or a two-tailed test (G). See also Figures S1, S8, S9, and S10.

### Pharmacologic EM inhibition

Finally, we tested the clinical relevance of our genetic findings and determined whether 6-Diazo-5-oxo-L-norleucine (DON), a broad-spectrum glutamine metabolism inhibitor investigated in myeloid leukemia^16^, could serve to inhibit EM-activated HSPCs. We combined 5FU treatment with DON, starting inhibition on D6 post-5FU (**Figure 6A**). Since continuous daily treatment impaired regeneration and led to 100% mortality within 14 days post-5FU, we limited DON treatment to 4 injections from D6 to D9 after 5FU injection. DON treatment during this window of HSPC activation led to delayed recovery of WBC counts followed by flagrant recovery of neutrophil output above that of the vehicle-treated mice (**Figure 6B**). BM analyses showed impaired expansion of HSCs and MPP3s during DON treatment at D8 post-5FU followed by delayed recovery of GMP after treatment cessation at D10 post-5FU (**Figure S11A**). Correspondingly, we observed a lack of GMP patches at D10, followed by the emergence of hypertrophic GMP clusters as myelopoiesis rebounded upon drug cessation at D12 post-5FU (**Figure S11B**). Finally, *in vivo* treatment with dmKG also rescued the delayed recovery of peripheral neutrophils post-5FU observed with DON treatment (**Figure S11C**). These results confirm our genetic results and demonstrate that pharmacological inhibition of glutaminolysis is sufficient to prevent HSPC-driven myeloid expansion, though sustained treatment will likely be required to prevent rebound and maintain the effect.

**Figure 6.**
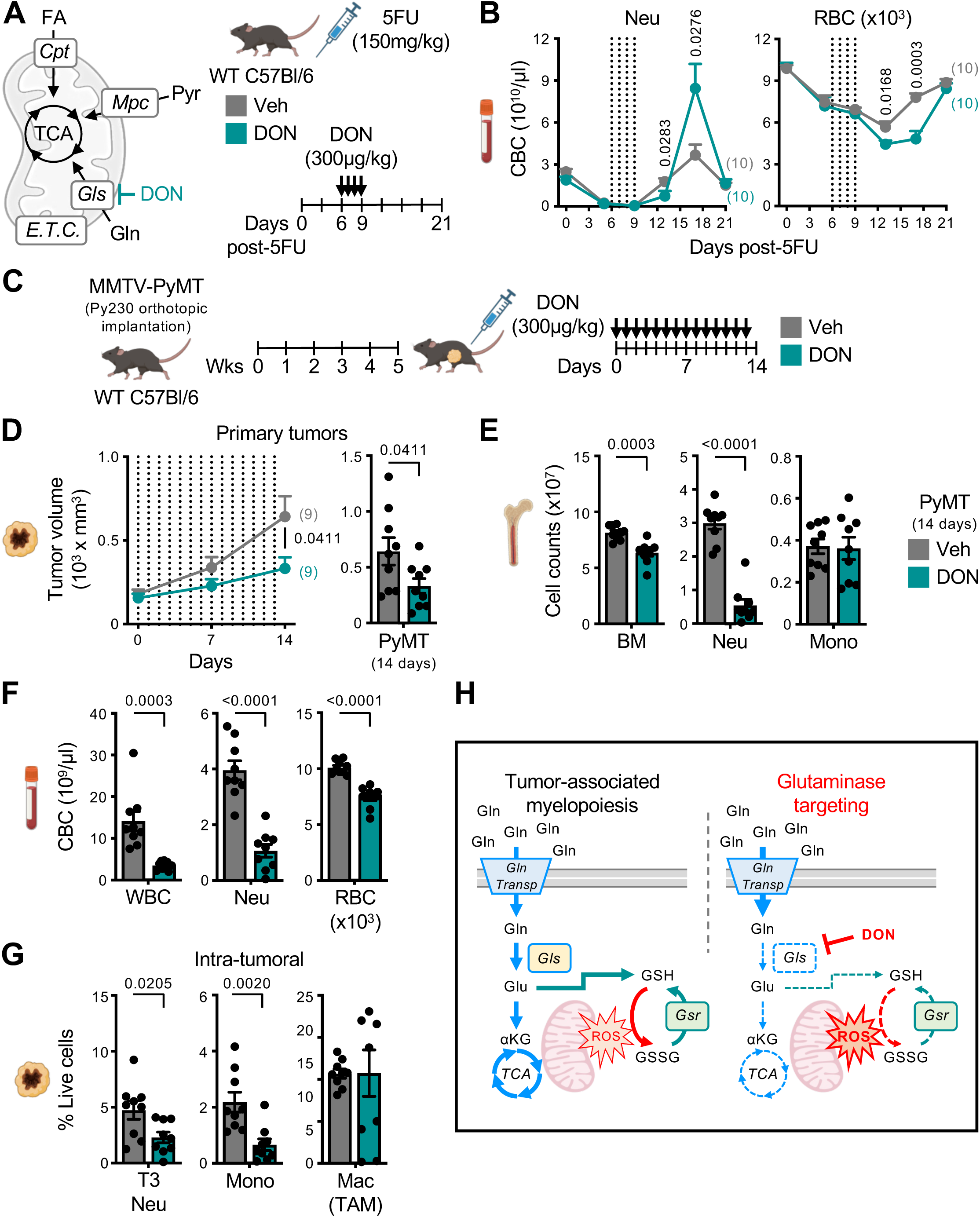
Pharmacologic inhibition of glutamine metabolism impairs regenerative and tumor-associated myelopoiesis. (A-B) Regenerative response in mice treated with the glutamine metabolism inhibitor 6-Diazo-5-oxo-L-norleucine (DON) or PBS vehicle (Veh) control: (A) treatment scheme with daily DON injections of days 6 to 9 post-5FU treatment; and (B) quantification of peripheral blood neutrophils and RBCs (n=10; 3 independent experiments). (C-G) Impact of pharmacological inhibition of glutaminolysis on breast cancer development in Veh and DON-treated WT mice (n=9; 1 independent experiments): (C) breast cancer model with daily DON injections 5 weeks after orthotopic implantation of MMTV-PyMT cell line (Py230, 1×106 cells); (D) breast tumor growth over time measured by external palpation (left), with tumor volume after 14 days of DON treatment; and quantification of (E) BM, (F) peripheral blood and (G) intratumoral myeloid populations after 14 days of DON treatment. (H) Model of glutaminase targeting to impair glutamine-driven OXPHOS and redox balance in myeloid progenitors as a novel therapeutic strategy to suppress tumor-associated myelopoiesis and neutrophil overproduction in breast cancer. Data are means ± S.E.M.; circles represent individual mice; *P. values* were obtained by an unpaired t-test. See also Figure S1, and S11.

We next evaluated the efficacy of DON treatment in tumor-bearing mice (**Figure 6C**). Strikingly, daily treatment with DON led to a significant reduction in tumor growth associated with a robust depletion of the BM neutrophil reservoir and decreased circulating neutrophils, as well as mild anemia in tumor-bearing WT mice (**Figure 6D to 6F**). This resulted in a decrease in both intratumoral neutrophils and monocytes, with no effect on tumor-associated macrophages (**Figure 6G**). Collectively, these data confirm our genetic findings and demonstrate that tumor-associated myelopoiesis exploits both EM-specific metabolic adaptation and lineage-specific metabolic dependency in myeloid progenitors (**Figure 6H**). Together, they establish the potential of EM HSPC glutamine addiction as a therapeutic target to alter the tumor microenvironment and slow tumor growth by starving the tumor of pro-angiogenic/pro-tumor neutrophils.

## DISCUSSION

Maladaptive EM pathway activation has long been considered a driver of disease progression; however, targeting myeloid overproduction without compromising normal hematopoiesis and the innate immune system has so far proved intractable^1^. This is primarily due to our limited understanding of the core mechanisms distinguishing steady state myelopoiesis from EM. Here, we identify the central regulatory axis of metabolic adaptation during EM, where hyperactivation of Myc drives increased mitochondrial biogenesis and OXPHOS, creating a critical dependence on glutaminolysis to fuel the TCA cycle. This dependence on glutaminolysis is highly specific to EM, as genetic or pharmacological glutaminase deficiency has a minimal impact on steady state myelopoiesis, which depends more heavily on aerobic glycolysis^10^. In addition, we found that the biosynthesis of glutamine, rather than its catabolism to fuel OXPHOS, is a key survival mechanism allowing quiescent HSCs to persist and maintain their proteostasis independently of exogenous glutamine, which also render them insensitive to the effects of glutaminolysis inhibition. In this context, it is likely that AML cells are co-opting the de novo glutamine biosynthesis capacity of HSCs for their own survival under conditions of glutamine deprivation^38^. Altogether, we show that EM-activated HSPCs resemble neoplastic tissues, where Myc-driven “glutamine addiction” has long been described as a key feature of cancer cell metabolism^19^. In fact, glutamine metabolism, and glutaminolysis specifically, have been identified as therapeutic targets in malignant hematopoiesis^16,39–44^. Rather than being cancer-specific features, we establish that these metabolic adaptations represent a shared biology allowing for demand-adapted myeloid production in normal hematopoiesis, which is hijacked for AML proliferation in disease conditions. Our results therefore place glutamine and its metabolism, its biosynthesis and catabolism, as central regulators of HSC behavior and hematopoietic plasticity.

Targeting EM-specific metabolic adaptation in HSPCs represents a novel immunotherapeutic strategy to suppress pathogenic neutrophil production and restructure the tumor microenvironment in breast cancer. While an early clinical trial investigating the efficacy of glutaminase inhibition in advanced metastatic triple-negative breast cancer (NCT03057600) was unsuccessful, we believe this reflects an unfortunate patient selection focused on cancer cell intrinsic glutamine metabolism, without consideration of the impact on EM-driven myelopoiesis. Further clinical evaluation of glutaminase inhibitors should instead focus on cancer subtypes in which neutrophil infiltration and/or the neutrophil-to-lymphocyte ratio (NLR)^45^ is associated with a poor prognosis, such as in ER+ breast cancer patients^46^, or is predictive of poor treatment response^47,48^. Our observation of myelopoietic rebound also indicates that sustained treatment will be a critical requirement for the utilization of glutamine metabolism inhibitors to target EM, and preclinical studies suggest they should be further tested in combination with immune checkpoint blockade^49–51^. Collectively, our work supports revisiting glutamine inhibition for breast cancer treatment with an understanding of its activity as a myeloid targeting agent. It also highlights the therapeutic potential of repurposing anti-leukemic agents as novel immunotherapeutic agents to modulate the innate immune system in solid cancers. Ultimately, we believe the proof-of-concept study presented here, while focused on solid cancer-associated myelopoiesis, will be relevant to other chronic inflammatory conditions where myeloid overproduction drives disease pathogenesis.

### Limitations of the study

Several limitations of this study should be considered when interpreting our findings. These studies have been carried out using mouse models, and while the metabolic dependencies we describe here are likely conserved, direct validation in human HSCPs will be required to confirm the translational relevance of our findings. While we have confirmed our results in several models of regenerative and maladaptive EM (*e.g*., 5FU, G-CSF, Py230) and validated the therapeutic potential of metabolic targeting of EM in a mouse model of breast cancer as a proof of concept, using additional breast cancer modes as well as other solid cancer models and inflammatory disease conditions will be required to evaluate the full clinical potential of HSPC glutaminase targeting. Additionally, while DON has favorable potency and biological activity, and can mimic the effect of hematopoietic-specific genetic glutaminase deletion, it can also inhibit other glutamine-dependent metabolic processes^16^. Testing additional more specific glutaminase inhibitor suc as the now discountinued CB-839 (Telaglenastat)^39,43^ will be warranted. Since we tested the requirement for Myc in driving EM using an haploinsufficiency approach, as complete Myc deletion is lethal^25^, it is possible that we underestimated the full contribution of Myc to the metabolic reprogramming of EM. Finally, single-cell metabolic flux estimation using scFEA provides a computational approximation of metabolic activity inferred from transcriptomic data, and while we validate key findings with direct functional measurements, including Seahorse analysis and stable isotope tracing, scFEA predictions should be interpreted as supportive rather than definitive evidence of flux changes.

## RESOURCE AVAILABILITY

All newly generated datasets supporting this study have been deposited in the Gene Expression Omnibus (GEO) under accession numbers GSE318312 and GSE298048. The M. musculus genome GRCm38.p4 is available from the National Center for Biotechnology Information (https://www.ncbi.nlm.nih.gov/assembly/GCF_000001635.24/). The chip annotation MSigDB.v7.2.chip is available from the Broad Institute (https://software.broadinstitute.org/cancer/software/gsea/wiki/index.php/MSigDB_v7.2_Release_Notes). Gene sets used for GSEA h.all.v7.2.symbols.gmt are available from the Broad Institute (https://data.broadinstitute.org/gsea-msigdb/msigdb/release/7.2/). All code and packages used to support the findings of this manuscript are either publicly available or available upon reasonable request. Correspondence and requests for materials should be addressed to E.P.

## ACKNOWLEDGEMENTS

We thank Bradley Allen, J. Perez Bruno and Brandon Wright (CUIMC) for broad technical assistance, Aimee A. Flores and Carlos Galvan (UCLA) for their help with the metabolomic studies, Dr. Ross Levine (MSKCC) for the *Scl-Cre^ER^* mice, Dr. Stephen Rayport (CUIMC/NYSPI) for the *Gls* floxed mice, Dr. Michel Nussenzweig (Rockefeller University) for the *Myc* floxed mice, M. Kissner and team members for excellent operation of the CSCI Flow Cytometry Core (CUIMC), Erin Bush and team members for managing the Single Cell Analysis Core (CUIMC), Heather Christofk and members of her laboratory for metabolomics analyses (UCLA), and all members of the Passegué laboratory for critical insights and suggestions. O.C.O was supported by Cancer Research Institute (CRI3617 Margaret Dammann Eisner Fellow), M.A.P. by NIH TL1DK136048, and J.W.S. by EMBO ALTF-2021-196 and Damon Runyon Cancer Research Foundation (William Raveis Charitable Fund Fellow) DRG-2493-23. This work was funded by NIH R01 AR084245 to W.E.L. and NIH R01 CA255342 and R35 HL135763 to E.P. and supported in part through the NIH/NCI Cancer Center Support Grant P30 CA013696 to CUIMC and NIH UL1 TR001873 to the Genomics and High Throughput Screening Shared Resource at CUIMC.

## AUTHOR CONTRIBUTIONS

Conceptualization: OCO, EP; Methodology: OCO, MAP, RZ, JWS, WEL; Investigation: OCO, MAP, RZ, JWS; Visualization: OCO, RZ, JWS; Funding acquisition: EP; Project administration: OCO, EP, WEL; Supervision: OCO, EP, WEL; Writing – original draft: OCO, EP; Writing – review & editing: OCO, EP, JWS, WEL.

## METHODS

### Resource Availability

#### Lead Contact

Further information and requests for resources and reagents should be directed to and will be fulfilled by the lead contact, Emmanuelle Passegué

#### Materials Availability

This study did not generate new unique reagents. Mouse lines used in this study are available from the Jackson Laboratory (see Key Resources Table) or upon request from the lead contact with a completed materials transfer agreement.

#### Data and code availability

- Single-cell RNA sequencing and Multiome data generated in this study have been deposited at GEO under accession numbers GSE318312 and GSE298048 and are publicly available as of the date of publication. Accession numbers are also listed in the Key Resources Table.
- This paper does not report original code. All packages used to support the findings of this manuscript are publicly available (see Key Resources Table for URLs). Any custom analysis scripts are available from the lead contact upon reasonable request.
- Any additional information required to reanalyze the data reported in this paper is available from the lead contact upon request.

### Experimental Model and Study Participant Details

#### Mice

All animal experiments were conducted at the Columbia University Irving Medical Center (CUIMC) in accordance with approved Institutional Animal Care and Use Committee protocols, and in compliance with all relevant ethical regulations. Wild type (WT) donor C57BL/6J (CD45.2) and recipient B6.SJL-Ptprca Pepcb/BoyJ (CD45.1) mice of both sexes were purchased from the Jackson Laboratory and bred in-house to generate all experimental animals. *MMTV-PyMT* mice^52^ (FVB/N-Tg(MMTV-PyVT)634Mul/J), *Myc-eGfp* mice^22^ (Myctm1.1Dlev/J), and *MRP8-Cre-ires-Gfp* mice^33^ (B6.Cg-Tg(S100A8-cre,-EGFP)1Ilw/J) were purchased from the Jackson Laboratory. *Scl-Cre^ER^*mice^23^ (C57BL/6-Tg(Tal1-cre/ERT)42-056Jrg/J) were obtained from Dr. Ross Levine (MSKCC), *Gls* floxed mice^32^ (Glstm2.1Sray/J) from Dr. Stephen Rayport (Columbia University and NYSPI), and *Myc* floxed mice^24^ (B6.129S6-Myctm2Fwa/Mmjax) from Dr. Michel Nussenzweig (Rockefeller University). For Cre-mediated deletion, ∼8-week-old control and *Scl-Cre^ER^*littermates were treated three times, 3 days apart, with 4 mg tamoxifen in 200 μl corn oil by oral gavage and were used for *in vivo* or *in vitro* experiments 4 weeks after the initial tamoxifen treatment. Both male and female animals were used in most experiments. Only female mice were used for breast cancer orthotopic implantation experiments. No specific randomization or blinding protocol was employed regarding the identity of experimental animals. Recipient mice for transplantation assays were 8–12 weeks old at the time of irradiation and cell transfer. Animal facilities were maintained at 22 ± 2 °C and 50 ± 10% relative humidity on a 12–12 hours light–dark cycle, and mice were given ad libitum access to Purina LabDiet Rodent Feed and acidified water. Mice were euthanized by CO asphyxiation followed by cervical dislocation.

#### Cell line

The syngeneic MMTV-PyMT breast cancer cell line Py230^35^ (ATCC CRL-3279) was used for orthotopic tumor implantation experiments. Cells were implanted at 1×10 cells per mouse in 100 μl of 50% Matrigel Growth Factor Reduced (GFR) Basement Membrane Matrix (Corning, Cat#354230) in PBS into the mammary fat pad of 8–10-week-old female C57BL/6J mice.

### Method Details

#### In vivo treatments

5-fluorouracil (5FU) and G-CSF treatments were performed as previously described^21^. In brief, mice were injected i.p. with either a single 150 mg/kg dose of 5FU (Sigma-Aldrich, Cat#F6627) in PBS or daily for 6 days with 5 μg human G-CSF (Neupogen, Amgen) in PBS. Treatments with 6-Diazo-5-oxo-L-norleucine (DON) were performed as previously described^16^. In brief, mice were injected i.p. with 300 μg/kg DON (Sigma-Aldrich, Cat#D2141) in PBS. During 5FU treatment, DON was injected on days 6, 7, 8, and 9. In the context of breast cancer, DON was injected daily starting 5 weeks after Py230 orthotopic implantation. Treatments with dimethyl α-ketoglutarate (dmKG) were performed as previously described^53^. In brief, on days 6, 7, 8, and 9 following 5FU treatment, mice were injected i.p. with 300 mg/kg of dimethyl α-ketoglutarate (Sigma-Aldrich, Cat#349631) in PBS.

#### Transplantation

For transplantation experiments, CD45.1 recipient mice were irradiated with 10.5 Gy delivered in split doses 3-4 hours apart using an X-ray irradiator (MultiRad225, Precision X-Ray). For CD45.2 HSCs cultured in self-renewal conditions, the content of each well seeded with 1,000 HSCs was recovered after 72 hours of culture and injected retro-orbitally alongside 600,000 Sca-1-depleted CD45.1 helper BM cells into 4 recipient mice. All recipient mice received water containing polymyxin (1,000 U/ml, Sigma-Aldrich Cat#P4932) and neomycin (1.2 mg/ml, Sigma-Aldrich Cat#N1876) for 4 weeks following transplantation. Peripheral blood was collected with capillary tubes under isoflurane anesthesia by retro-orbital bleeding and either dispensed into EDTA-coated tubes for complete blood count (CBC) analyses using a Genesis hematology system (Oxford Science) or collected directly into 1 mM EDTA-containing ACK (150 mM NH_4_Cl and 10 mM KHCO_3_) red blood cell (RBC) lysis buffer for flow cytometry staining.

#### Flow cytometry of bone marrow hematopoietic cells

BM cells were obtained by crushing 2 pelvic bones, 2 humeri, 2 tibias and 1 femur in staining medium composed of HBSS (Gibco, Cat#14175103) containing 2% heat-inactivated FBS (Gibco, Cat#16140-071). RBCs were removed by lysis with ACK buffer, and single-cell suspensions were purified on a Ficoll gradient (Histopaque 1119, Sigma-Aldrich). BM cells were pre-enriched for c-Kit^+^ cells using c-Kit microbeads (Miltenyi Biotec, Cat#130-091-224) and an AutoMACS Pro separator. For HSPC analyses and isolation of LK (Lin^□^/c-Kit^+^), LSK (Lin^□^/Sca-1^+^/c-Kit^+^), HSC (LSK/Flk2^□^/CD48^□^/CD150^+^), MPP3 (LSK/Flk2^□^/CD48^+^/CD150^□^), and GMP (LK/Sca-1^□^/CD34^+^/CD16/32^+^) BM populations, cells were incubated with the following antibodies: c-Kit-APC-Cy7 (BioLegend, Cat#105826; 1:800), Sca-1-BV421 (BioLegend, Cat#108128; 1:400), CD150-BV650 (BioLegend, Cat#115931; 1:200), CD48-AF700 (BioLegend, Cat#103426; 1:400), Flk2-PE (eBioscience, Cat#12-1351-82; 1:100), CD34-FITC (eBioscience, Cat#11-0341-85; 1:25), CD16/32-PE/Cy7 (BioLegend, Cat#101318; 1:800), Ly-6C-APC (BioLegend, Cat#128016; 1:600), and CD115-BV711 (BioLegend, Cat#135515; 1:400) along with lineage antibodies in PE/Cy5 (Gr-1 [eBioscience, Cat#15-5931-82; 1:800], CD11b [Invitrogen, Cat#15-0112-82; 1:800], B220 [Invitrogen, Cat#15-0452-82; 1:800], CD19 [BioLegend, Cat#115510; 1:800], CD5 [BioLegend, Cat#100610; 1:800], CD3 [Invitrogen, Cat#15-0031-83; 1:400], CD4 [Invitrogen, Cat#15-0041-82; 1:800], CD8a [Invitrogen, Cat#15-0081-82; 1:800], Ter119 [Invitrogen, Cat#15-5921-83; 1:400]). For some analyses using transgenic reporter mice or fluorescent molecular probes, CD34-biotin (BioLegend, Cat#119304; 1:50) or Flt3-biotin (BioLegend, Cat#135308; 1:100) was substituted, combined with Streptavidin-BV605 (BioLegend, Cat#405229; 1:400). For dye analyses, after surface staining, cells were resuspended in 40 nM TMRE (Enzo Life Sciences, Cat#ENZ-52309) or 800 nM ThiolTracker Violet (ThermoFisher, Cat#T10095) in HBSS and incubated for 30 min at 37°C. For myeloid profiling and isolation of BM neutrophils (Lin^□^/CD11b^+^/Ly6C^+^/Ly6G^+^), unfractionated BM cells were stained with c-Kit-BV785 (BioLegend, Cat#105841; 1:800), CD115-PE (BioLegend, Cat#135505; 1:400), CD11b-PE/Cy7 (Invitrogen, Cat#25-0112-82; 1:800), CD101-APC (Invitrogen, Cat#17-1011-82; 1:800), Ly-6G-AF700 (BioLegend, Cat#127622; 1:400), and Ly-6C-APC/Cy7 (BioLegend, Cat#128026; 1:600) along with lineage antibodies in PE/Cy5. For chimerism analyses, peripheral blood cells were stained with CD45.1-PE (Invitrogen, Cat#12-0453-83; 1:400), CD45.2-FITC (Invitrogen, Cat#111-0454-85; 1:400), Gr-1-eF450 (Invitrogen, Cat#48-5931-82; 1:800), CD11b-PE/Cy7 (Invitrogen, Cat#25-0112-82; 1:800), CD8-AF700 (BD, Cat#557959; 1:400), CD4-BV510 (BioLegend, Cat#100553; 1:400), CD3-APC (BioLegend, Cat#100236; 1:100), B220-APC/Cy7 (BioLegend, Cat#103224; 1:800). Stained cells were finally resuspended in staining media containing 1 μg/ml propidium iodide for dead cell exclusion. HSPC cell isolations were performed on a FACSAria II SORP (BD Biosciences) using double sorting for purity. Cell analyses were performed on a NovoCyte Quanteon or Penteon analyzer (Agilent Technologies). Data collection was done using BD FACSDiva (v9) or Agilent NovoExpress (v1.6.2), and data analysis was performed using FlowJo (v10).

#### Flow cytometry of intratumoral immune cells

After euthanasia, mammary tumors were excised and a <1 cm³ piece was placed in 5 ml digestion buffer containing 300 U/ml Collagenase (Fisher Scientific, Cat#NC9096128), 100 U/ml Hyaluronidase (Fisher Scientific, Cat#NC9928421), 2 μg/ml DNase I (Sigma-Aldrich, Cat#10104159001) in HBSS/2% FBS. Tumor fragments were minced to <50 mm³ and digested on a gentleMACS Octo Dissociator using the “37C_m_TDK_2” program. Tumor digests were filtered through 45 μm mesh strainers, ACK-treated to remove red blood cells, and stained for flow cytometry. For tumor myeloid profiling, tumor digests were stained with CD45.2-BV421 (Invitrogen, Cat#404-0454-80; 1:400), CD11b-PE/Cy7 (Invitrogen, Cat#25-0112-82; 1:800), Ly-6G-AF700 (BioLegend, Cat#127622; 1:400), Ly-6C-APC/Cy7 (BioLegend, Cat#128026; 1:600), CD101-APC (Invitrogen, Cat#17-1011-82; 1:800), CD115-BV711 (BioLegend, Cat#135515; 1:400), DcTRAIL-R1-PE (BioLegend, Cat#133804; 1:200) along with lineage antibodies in PE/Cy5 (B220, CD3, NK-1.1, Ter119, EpCAM). For tumor lymphoid profiling, tumor digests were stained with CD45.2-BV421 (1:400), CD3-APC (BioLegend, Cat#100236; 1:100), CD4-BV510 (BioLegend, Cat#100553; 1:400), CD8-AF700 (BD, Cat#557959; 1:400), NK-1.1-PE (BioLegend, Cat#108708; 1:800) along with lineage antibodies in PE/Cy5 (Gr-1, CD11b, Ter119, EpCAM). Stained cells were finally resuspended in staining media containing 1 μg/ml propidium and analyzed on a NovoCyte Quanteon or Penteon.

#### In vitro liquid culture assays

All cultures were performed at 37°C in a 5% CO water jacket incubator in normoxia. For PVA self-renewal culture conditions, HSCs or MPP3s (1,000 cells/well) were deposited in 96-well flat-bottom plates (Falcon, Cat#353071) pre-coated with 1 μg/ml bovine fibronectin (Sigma-Aldrich, Cat#F1141) for 30 min at 37°C, into 150 μl of PVA media^54^: DMEM:F12 (Thermo Scientific, Cat#21331020) containing 100 U/ml penicillin and 100 μg/ml streptomycin (Thermo Scientific, Cat#15140122), 10 mM HEPES (Cat#15-630-080), 1 mg/ml polyvinyl alcohol (Sigma-Aldrich, Cat#363146), ITS-X (Fisher Scientific, Cat#51500056), 10 ng/ml SCF (PeproTech, Cat#250-03), and 100 ng/ml TPO (PeproTech, Cat#315-14A). Where indicated, the PVA media also contained 2 mM L-glutamine (Fisher Scientific, Cat#35-050-061), 100 μM methionine sulfoximine (MSO, Fisher Scientific, Cat#AC227202500), or 200 nM rapamycin (EMD Millipore, Cat#553210). Cells were counted after 72 hours of culture by removing 100 μl of medium and mixing with 50 μl of HBSS/2% FBS and 1 μg/ml propidium iodide before counting on a NovoCyte Penteon using the absolute count setting. Incorporation of o-propargyl-puromycin (OP-Puro) was quantified after 3 hours of PVA culture by adding 20 μM o-propargyl-puromycin in PVA medium, incubating for an additional 1 hour, fixing and staining the cells according to manufacturer instructions in the Click-IT Plus OPP Alexa Fluor 647 Kit (Invitrogen, Cat#C10458), and analyzing them by flow cytometry on a NovoCyte Penteon. For myeloid expansion culture conditions, LSKs (100 cells/well) or GMPs (500 cells/well) were deposited in 96-well suspension culture plates (Greiner Bio-One, Cat#655-185) into 150 μl of myeloid expansion media: DMEM:F12 containing 5% FBS, 100 U/ml penicillin and 100 μg/ml streptomycin, 2 mM L-glutamine, and the following cytokines (all from PeproTech): 10 ng/ml IL-3, 10 ng/ml GM-CSF, 25 ng/ml SCF, 25 ng/ml IL-11, 25 ng/ml Flt3L, 25 ng/ml TPO, and 4 U/ml EPO. Where indicated, the culture media also contained 50 nM rotenone (Sigma-Aldrich, Cat#R8875), 5 μM etomoxir (Selleck Chemicals, Cat#S8244), 5 μM BPTES (bis-2-(5-phenylacetamido-1,2,4-thiadiazol-2-yl)ethyl sulfide, Sigma-Aldrich, Cat#SML0601), 5 μM UK5099 (Thermo Scientific, Cat#41-861-0), 400 μM dmKG (Sigma-Aldrich, Cat#349631), or 400 μM NAC (n-acetyl-L-cysteine, Sigma-Aldrich, Cat#A7250). Cells were counted after 72 hours of culture by removing 100 μl of medium and mixing with 50 μl of HBSS/2% FBS and 1 μg/ml propidium iodide before counting on a NovoCyte Penteon using the absolute count setting.

#### Metabolomics

For stable isotope tracing, mice were intraperitoneally injected with 300 mg/kg ¹³C L-glutamine (Cambridge Isotope Laboratories, Cat#CLM-1822-H-PK) in PBS 1 hour prior to euthanasia. LK, LSK, or GMP populations (150,000 cells) were sorted directly into 80% LC/MS grade methanol (Fisher Scientific, Cat#A456-1) and 10 mM trifluoromethanesulfonate (Sigma-Aldrich, Cat#372331) in LC/MS grade H O (Fisher Scientific, Cat#W6-4). After rigorous mixing, the suspension was pelleted by centrifugation (18,000g, 4°C). The supernatant was transferred to a fresh tube, and metabolites were dried under vacuum centrifugation before being shipped to UCLA and stored at −80°C. Prior to analyses, samples were resuspended in 70% acetonitrile, and mass spectrometry was performed as previously described^55^.

#### Seahorse extracellular flux analysis

Oxygen consumption rates (OCR) were measured using either an 8-well Seahorse XF HS Mini Analyzer (for LSK and GMP analyses) or a 96-well Seahorse Bioanalyzer XF96 (for neutrophil analyses) according to the manufacturer’s instructions (Agilent Technologies). In brief, 35,000 LSK or GMP cells and 200,000 neutrophils were sorted into 1.4 ml tubes, rested on ice for 1 hour (LSK/GMP) or 2 hours (neutrophils), and spun down at 400g for 5 min. Supernatant was aspirated to 20 μl and cells were resuspended with 180 μl of XF base medium (Agilent, Cat#103576-100) containing 10 mM glucose, 2 mM L-glutamine, 0.075% 1N NaOH, and for LSK/GMP analyses the cytokines of the myeloid expansion media. Cells were transferred into XF HS PDL Miniplates (LSK/GMP) or XF96 Cell Culture Microplates (neutrophils) and left to equilibrate for 1 hour at 37°C in a non-CO incubator. For mitochondrial stress tests, 10X injection dilutions were prepared for oligomycin A (final 5 μM; Sigma-Aldrich, Cat#75351), FCCP (final 2 μM; Sigma-Aldrich, Cat#C2920), and rotenone/antimycin A (final 1 μM/1 μM; Sigma-Aldrich, Cat#R8875/Cat#A8674). For oxidative burst tests^56^, 10X injection dilutions were prepared for PMA (final 5 μM; Sigma-Aldrich, Cat#P8139) and 2-DG (final 100 mM; Sigma-Aldrich, Cat#D3179).

#### BM section immunofluorescent imaging

Whole femurs were dissected, placed in plastic mounts containing OCT, and immediately frozen at −80°C. Sections of 7 μm thickness were cut using a Leica CM3050S cryostat and transferred to glass slides using a CryoJane tape transfer system (Leica). Slides were dried for 20 min at RT, then fixed in acetone at −20°C for 10 min. A circle was drawn around tissue sections using a PAP pen before washing three times with PBS. Slides were incubated for 1 hour at RT in 10% goat serum (Fisher Scientific, Cat#16-210-064) in PBS, washed, then stained with rat anti-mouse CD117 primary antibody (BioLegend, Cat#135101; 1:25 dilution) in blocking media for 2 hours at RT. After washing, slides were stained with goat anti-rat IgG Cy3 secondary antibody (Jackson ImmunoResearch, Cat#112-165-167; 1:100 dilution) for 30 min at RT, washed, and incubated with rat IgG isotype control (Sigma-Aldrich, Cat#I8015; 1:50) for 30 minutes at RT. Slides were then incubated for 90 min at RT with a mix of directly conjugated antibodies: CD16/32-AF647 (BioLegend, Cat#101314; 1:100), Sca-1-AF488 (BioLegend, Cat#108116; 1:150), CD150-AF488 (BioLegend, Cat#115916; 1:150), CD11b-AF488 (BioLegend, Cat#101217; 1:200), Gr1-AF488 (BioLegend, Cat#108418; 1:200), B220-AF488 (BioLegend, Cat#103226; 1:200), CD3-AF488 (BioLegend, Cat#100210; 1:100) and DAPI (5 μg/ml). Slides were mounted with ProLong Glass Antifade Mountant (Thermo Fisher, Cat#P36980). Images were acquired with a Leica SP8 inverted confocal microscope using a 20X objective lens and Leica Application Suite X (LAS X). A z-stack of 1 μm was acquired across sections and final images were displayed as maximum intensity projections.

#### Tomm20 immunofluorescent imaging

Cells were directly sorted into a PAP pen circle on poly-lysine-coated slides (Sigma-Aldrich, Cat#P0425) for HSC/MPP3 populations (∼5–10,000 cells/slide) or fibronectin-coated slides for GMPs (∼10,000 cells/slide) and allowed to settle for 30 min at RT. For fibronectin coating, charged slides (Fisher Scientific, Cat#12-550-15) were coated with 10 μg/ml bovine fibronectin for 1 hour at RT and washed 3 times with PBS. Cells were fixed with 4% PFA (Electron Microscopy Sciences, Cat#15710) for 10 min, then blocked and permeabilized for 1 hour at RT in Block/Perm media containing 10% goat serum (Fisher Scientific, Cat#16-210-064), 1% BSA (Sigma-Aldrich, Cat#A7030), and 0.05% Triton X-100 (Sigma-Aldrich, Cat#93443) in PBS. Slides were stained with rabbit anti-Tomm20 primary antibody (Cell Signaling Technology, Cat#42406; 1:100) in Block/Perm media for 2 hours at RT, washed, and stained with donkey anti-rabbit IgG AF488 secondary antibody (Jackson ImmunoResearch, Cat#711-545-152; 1:400) and DAPI (5 μg/ml) for 1 hour at RT. Slides were mounted with ProLong Glass Antifade Mountant. Images were acquired with a Leica SP8 inverted confocal microscope using a 40X objective and LAS X. A z-stack of 1 μm was acquired and final images were displayed as sum-intensity projections, with signal quantification measured using ImageJ2 (v2.14).

### Quantitative RT-PCR

HSC, MPP3, or GMP populations (5,000 cells/tube) were sorted directly into DNA/RNA Shield (Fisher Scientific, Cat#50-125-1706) and RNA was extracted using a Quick-RNA MicroPrep Kit (Zymo Research, Cat#50-444-593). A High-Capacity cDNA Reverse Transcription Kit (Thermo Fisher Scientific, Cat#4368813) was used to generate cDNA libraries. Gene expression was assessed on a CFX Opus 96 Real-Time PCR System (Bio-Rad) using SsoFast EvaGreen Supermix with Low ROX (Bio-Rad, Cat#1725210) and the following primers: Gls-F (5’-TTCGCCCTCGGAGATCCTAC-3’), Gls-R (5’-CCAAGCTAGGTAACAGACCCT-3’), Glul-F (5’-TGAACAAAGGCATCAAGCAAATG-3’), Glul-R (5’-CAGTCCAGGGTACGGGTCTT-3’), Actb-F (5’-GACGGCCAGGTCATCACTATTG-3’), Actb-R (5’-AGGAAGGCTGGAAAAGAGCC-3’).

### 10X scRNA-seq analyses

For generating scRNA-seq datasets, 30,000–50,000 LK or LSK cells were sorted into HBSS with 2% FBS, rested on ice for 1 hour before pelleting at 350g for 5 min at 4°C. GEM generation and 3’ RNA library preparation were performed according to 10X Genomics protocol CG000315 Rev E, targeting 5,000 cell data recovery. RNA libraries were pooled, sequenced on an Illumina NovaSeq 6000 (2×150 bp, 2.5 billion reads), and aligned using Cell Ranger (v7.0.1) to mouse genome mm10. Library concentrations and fragment sizes were evaluated using Qubit dsDNA HS assay kit (ThermoFisher Scientific, Cat#Q33230) and TapeStation D5000 DNA ScreenTape (Agilent Technologies, Cat#5067-5589). Subsequent analysis was performed using Seurat v5^57,58^. Standard quality control filtering removed cells with low gene counts (nFeature_RNA <1,000), low unique RNA molecules (nCount_RNA <2,000), or >7.5% mitochondrial genes. Cell type annotation was performed using HemaScribe^17^, with LT-HSC and ST-HSC analyzed as a single HSC population. B-cell progenitors and lymphoid cells were removed for myeloid-focused analyses. A steady-state reference dataset was generated by combining LK and LSK datasets from unstimulated mice and integrated using *PrepSCTIntegration*, *FindIntegrationAnchors*, and *IntegrateData functions*. Reference objects were processed using *NormalizeData*, *FindVariableFeatures*, *ScaleData*, *RunPCA* (dims=1:50) and *RunUMAP* (dims=1:15). For perturbation datasets, samples were projected onto the reference UMAP using F*indTransferAnchors* (dims=1:30) and *MapQuery*. The metabolic enzyme list was derived from the Comprehensive Mammalian Metabolic Enzyme Database^59^. Gene sets for Myc-driven mitochondrial biogenesis targets, Hallmark Myc Targets, and Hallmark Oxidative Phosphorylation were sourced from MSigDB v7.2^60^ and gene set activity scores were assigned using *AddModuleScore*. Differential gene expression was measured using *FindMarkers* and gene set enrichment analysis was performed using GSEA v4.4.0^61^. For feature plots, single-cell pathway activity was scored using AUCell v1.32.0^62^ with KEGG metabolic pathways from msigdb v1.18.0. Single-cell metabolic flux estimation (scFEA v1.1.2)^18^ was performed using a custom R wrapper integrating scFEA’s Python implementation with Seurat objects, calculating flux values for 168 metabolic modules and 70 metabolite balance values.

### 10X Multiome analyses

For single-cell Multiome datasets, 30,000–50,000 LK or LSK cells were sorted into HBSS/2% FBS, rested on ice for 1 hour before pelleting at 350g for 5 min at 4°C. Pelleted cells were lysed in chilled lysis buffer (10 mM Tris-HCl pH 7.4, 10 mM NaCl, 3 mM MgCl2, 0.1% Tween-20, 0.1% NP-40 Substitute, 0.01% Digitonin, 1% BSA, 1 mM DTT, 1 U/ml RNase inhibitor) and incubated for 3 min on ice. Nuclei were washed, pelleted at 500g for 10 min at 4°C, and resuspended in diluted nuclei buffer (10X Genomics). Multiome GEM generation and library preparation was performed according to 10X Genomics protocol CG000338, targeting 5,000 cell data recovery. RNA data were processed and annotated as described for scRNA-seq. ATAC data were analyzed in Signac v1.14.0^63^ with the following QC parameters: ATAC count >1,000 and <100,000 per cell, nucleosome signal <2, TSS enrichment >1. Samples were integrated using *IntegrateEmbeddings*, and peaks were called for each dataset with MACS2 and combined into a merged peak file. After integration, PCA was performed with 30 components followed by UMAP using components 2–30. Differential peak accessibility was identified using *FindMarkers*, and motif enrichment was assessed using *FindMotifs*.

### Quantification and statistical analysis

All experiments were repeated as indicated; n indicates the number of independent biological replicates. Data are expressed as means ± standard error of the mean (S.E.M.) or ± standard deviation (S.D.) as indicated in figure legends. Statistical analyses were performed using GraphPad Prism (v9/v10). Data distribution was assumed to be normal but was not formally tested. Statistical tests used are indicated in the figure legends and include the unpaired two-tailed Student’s t-test, the Kruskal-Wallis test, the Kolmogorov-Smirnov test, and simple linear regression with a two-tailed test for slope significance. Exact n values, the number of independent experiments, and the specific statistical tests used for each experiment are reported in the figure legends. Sample sizes were predetermined based on our experience in mouse HSC biology, where a standard deviation of less than 10% is expected for most measurements. To show a difference of at least 10% with α = 0.05 and 80% power, 6–11 independent biological replicates are required per group. Larger numbers of replicates were used in experiments with high variability, such as drug treatments and transplantation assays. For transplantations, recipients with <1% overall chimerism were excluded from lineage quantification. Mice used for treatment and transplantation experiments were assigned to groups based on genotype, randomized by sex, and samples were alternated whenever possible. Data collection and analysis were not performed blind to the experimental conditions, except for blinded scoring of immunofluorescence images.

## SUPPLEMENTAL INFORMATION TITLES AND LEGENDS

**Document S1**. Figures S1–S11 with figure legends.

