## Supplemental Info for "Glutamine addiction is a therapeutic target to block emergency myelopoiesis"

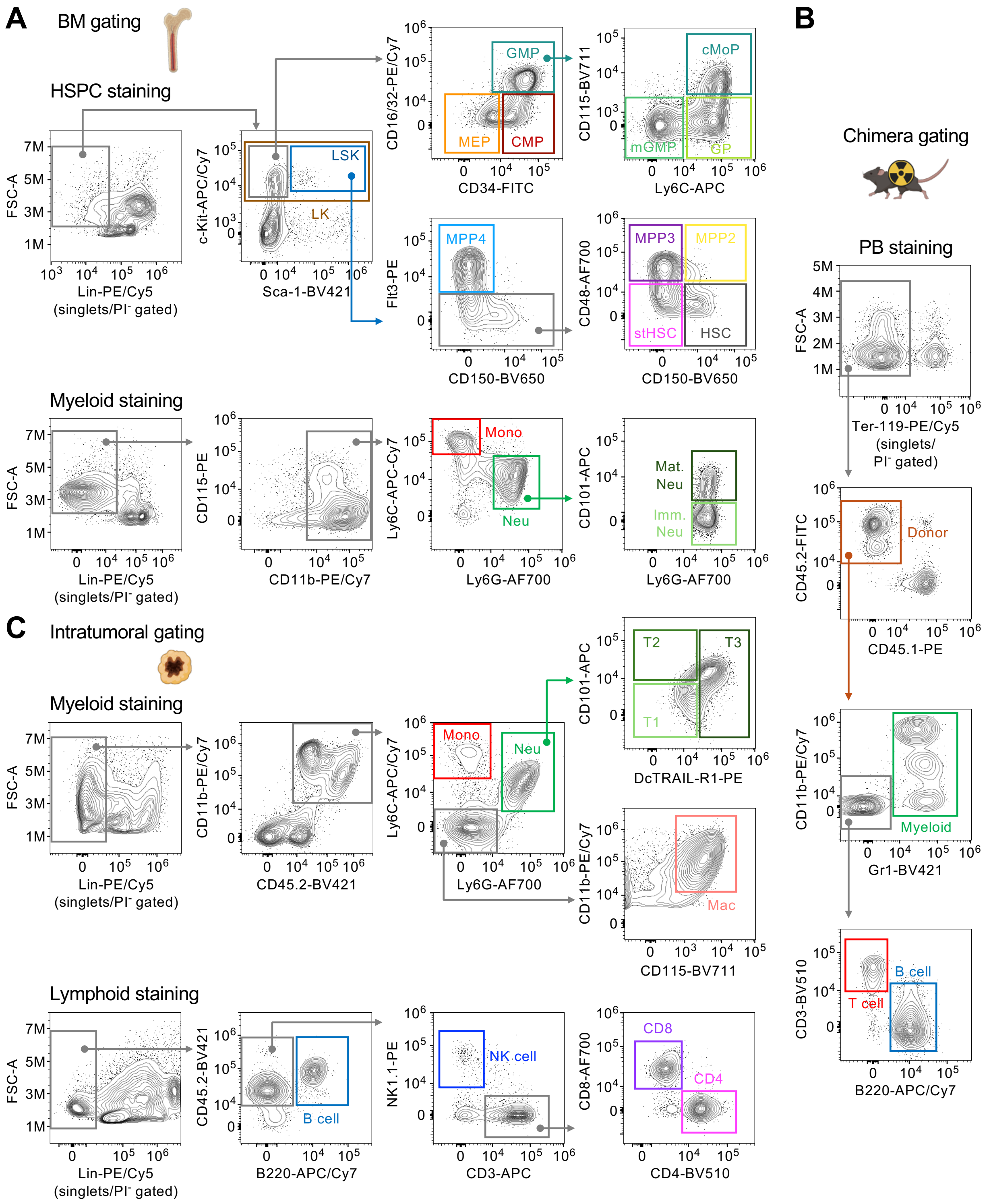


Figure S1 (Olson et al.)

**Figure S1. Examples of flow cytometric gating for identification of cell types, related to Figure 1, Figure 5, and Figure 6.**

(A) Flow cytometric gating for identification of bone marrow (BM) hematopoietic stem and progenitor cell (HSPC, top) and myeloid (bottom) populations. Lineage (Lin) includes Gr-1/CD11b/B220/CD19/CD3/CD5/CD4/CD8/Ter-119 for HSPC staining and B220/CD19/CD3/CD5/CD4/CD8/Ter-119 for myeloid staining. LK, $Lin^{+}$/c-$Kit^{+}$ cell; LSK, $Lin^{+}$/Sca-$1^{+}$/c-$Kit^{+}$ cell; HSC, hematopoietic stem cell; stHSC, short-term hematopoietic stem cell; MPP, multipotent progenitor; MEP, megakaryocyte/erythroid progenitor; CMP, common myeloid progenitor; GMP, granulocyte/macrophage progenitor; mGMP, multi-lineage granulocyte/macrophage progenitor; GP, granulocyte progenitor; cMoP, common monocyte progenitor; Mono, monocyte; Neu, neutrophil; Mat., mature; Imm., immature.

(B) Flow cytometric gating for identification of chimeric populations in the peripheral blood (PB).

(C) Flow cytometric gating for identification of intratumoral myeloid (top) and lymphoid (bottom) populations. Lineage (Lin) includes EpCAM/B220/CD3/NK1.1/Ter-119 for myeloid staining and EpCAM/CD11b/Gr-1/Ter-119 for lymphoid staining. Mac, macrophage; NK cell, natural killer cell.

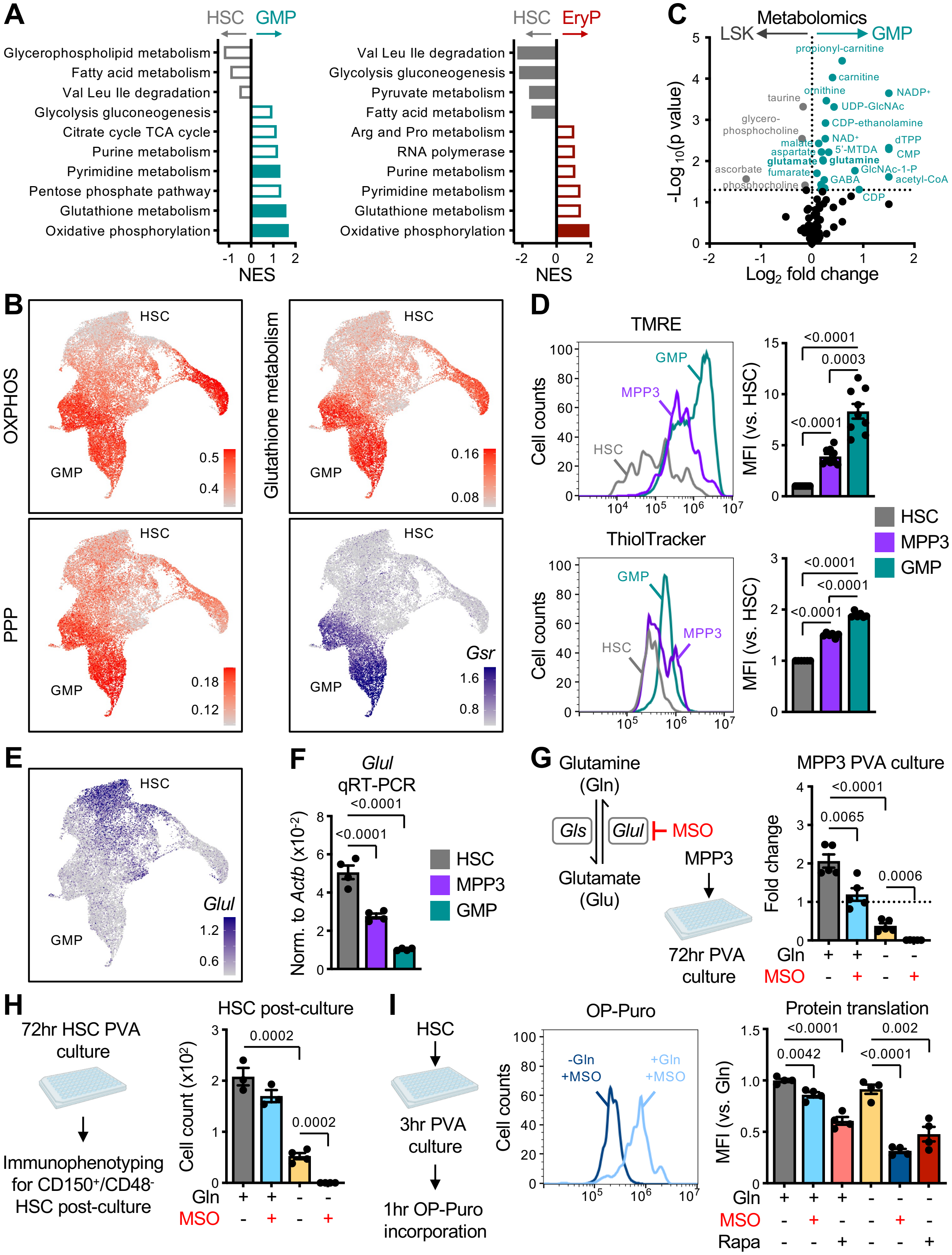


Figure S2 (Olson et al.)

**Figure S2. Hematopoietic differentiation is defined by lineage-rooted metabolic identities, related to Figure 1.**
(A) KEGG GSEA analysis of transcriptional changes as a function of myeloid (GMP vs. HSC, left) and erythroid (EryP vs. HSC, right) lineage commitment; open bars = non-significant; closed bars = significant ($p < 0.05$); NES, normalized enrichment score.
(B) Feature plots of lineage-specific KEGG metabolic pathway activity (red) and metabolic gene (blue). PPP, pentose phosphate pathway; Gsr, glutathione reductase.
(C) LC-MS metabolomic analysis of LSK and GMP (n=6; 2 independent experiments), with volcano plot of analyzed metabolites.
(D) Measurement of mitochondrial membrane potential (top) and reduced glutathione pool (bottom) in BM HSPCs by TMRE (n=9; 3 independent experiments) and ThiolTracker (n=6; 3 independent experiments), respectively.
(E) Feature plot of the HSC-specific metabolic gene. Glul, glutamine synthetase.
(F) Validation of Glul expression by qRT-PCR in isolated HSPCs (n=4; 2 independent experiments).
(G-I), Response to glutamine starvation following PVA culture with or without glutamine (±Gln) and the glutamine biosynthesis inhibitor methionine sulfoximine (±MSO): (G) MPP3 expansion rates after 72 hours PVA culture expressed as fold change of plated cells (n=5; 2 independent experiments); (H) immunophenotypic quantification of $CD150^{+}/CD48^{+}$ HSCs obtained after 72 hours HSC PVA culture (n=3-4; 2 independent experiments); and (I) quantification of protein translation measured by 1 hour OP-Puro incorporation following 3 hours HSC PVA culture (n=4; 3 independent experiments); Rapa, rapamycin.
Data are means ± S.E.M.; circles represent individual mice; *P. values* were obtained by the Kolmogorov-Smirnov test (A), and an unpaired t-test (C, F, G, H, I)..

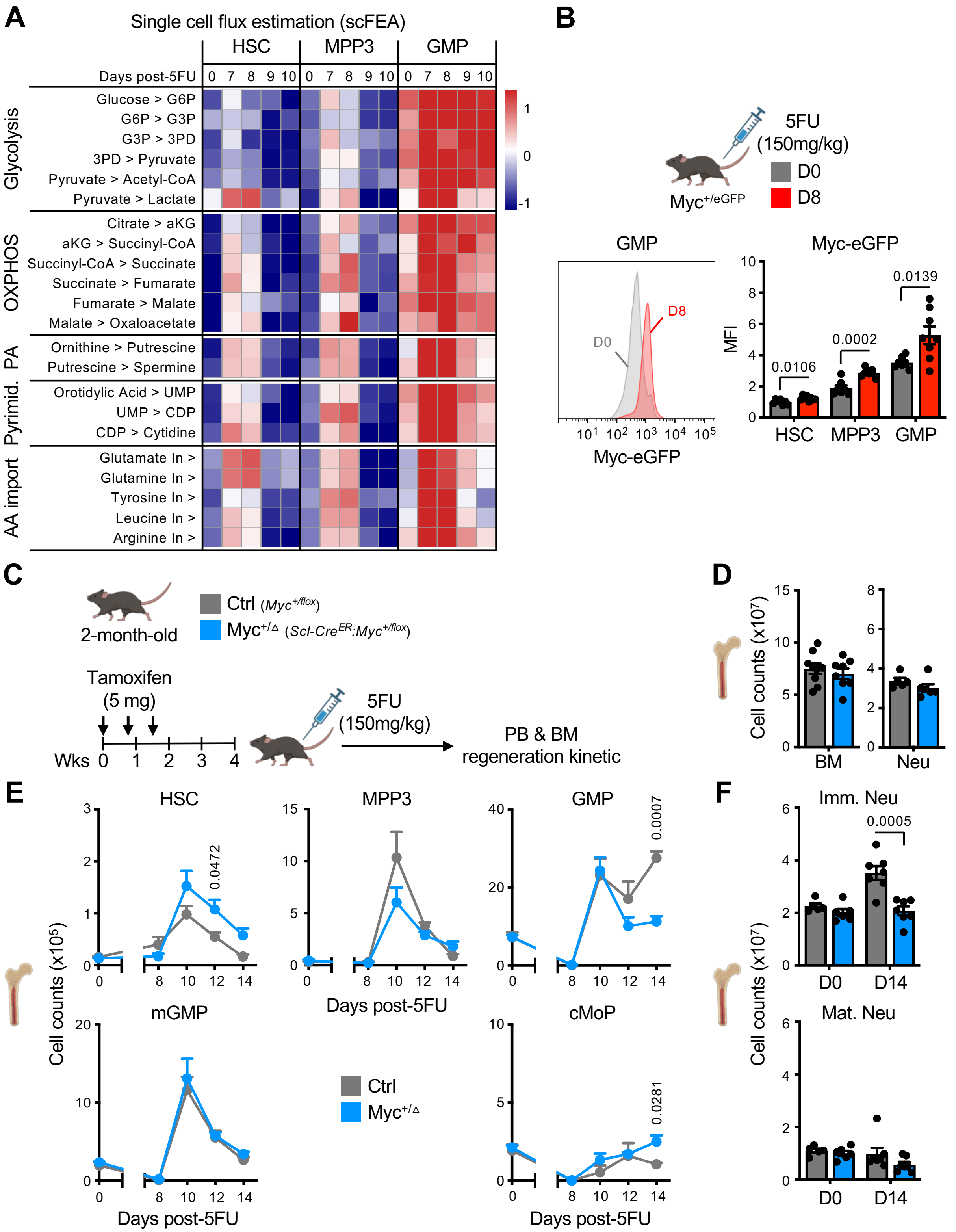


Figure S3 (Olson et al.)

**Figure S3. Myc hyperactivation drives the regeneration of hematopoietic progenitors, related to Figure 2.**

(A) scFEA analysis of indicated metabolic pathways in regenerative HSPCs post-5FU treatment. PA, polyamine metabolism; Pyrimid, pyrimidine nucleotide biosynthesis; AA import, amino acid import.

(B) Myc protein levels in regenerative HSPCs at day 0 (D0) and day 8 (D8) post-5FU isolated from a *Myc-eGfp* mouse reporter strain (n=7-8; 2 independent experiments). MFI, mean intensity fluorescence.

(C-F), Analysis of hematopoietic-specific Myc haplo-insufficient mice: (C) scheme of tamoxifen-mediated Myc inactivation (wks, weeks) and 5FU treatment in control (Ctrl) and hematopoietic-specific Myc-deleted ($Myc^{+/\Delta}$) mice; and quantification of (D) BM cellularity and neutrophils at steady state (n=5-7; 2 independent experiments), (E) BM HSPC post-5FU treatment (n=3-7 mice per time point and genotype; 3 independent experiments), and (F) BM $CD101^-$ immature neutrophils (Imm. Neu) and $CD101^+$ mature neutrophils (mat. Neu) at day 0 (D0) and day 14 (D14) post-5FU (n=5-7; 2 independent experiments).

Data are means ± S.E.M.; circles represent individual mice; *P. values* were obtained by an unpaired t-test.

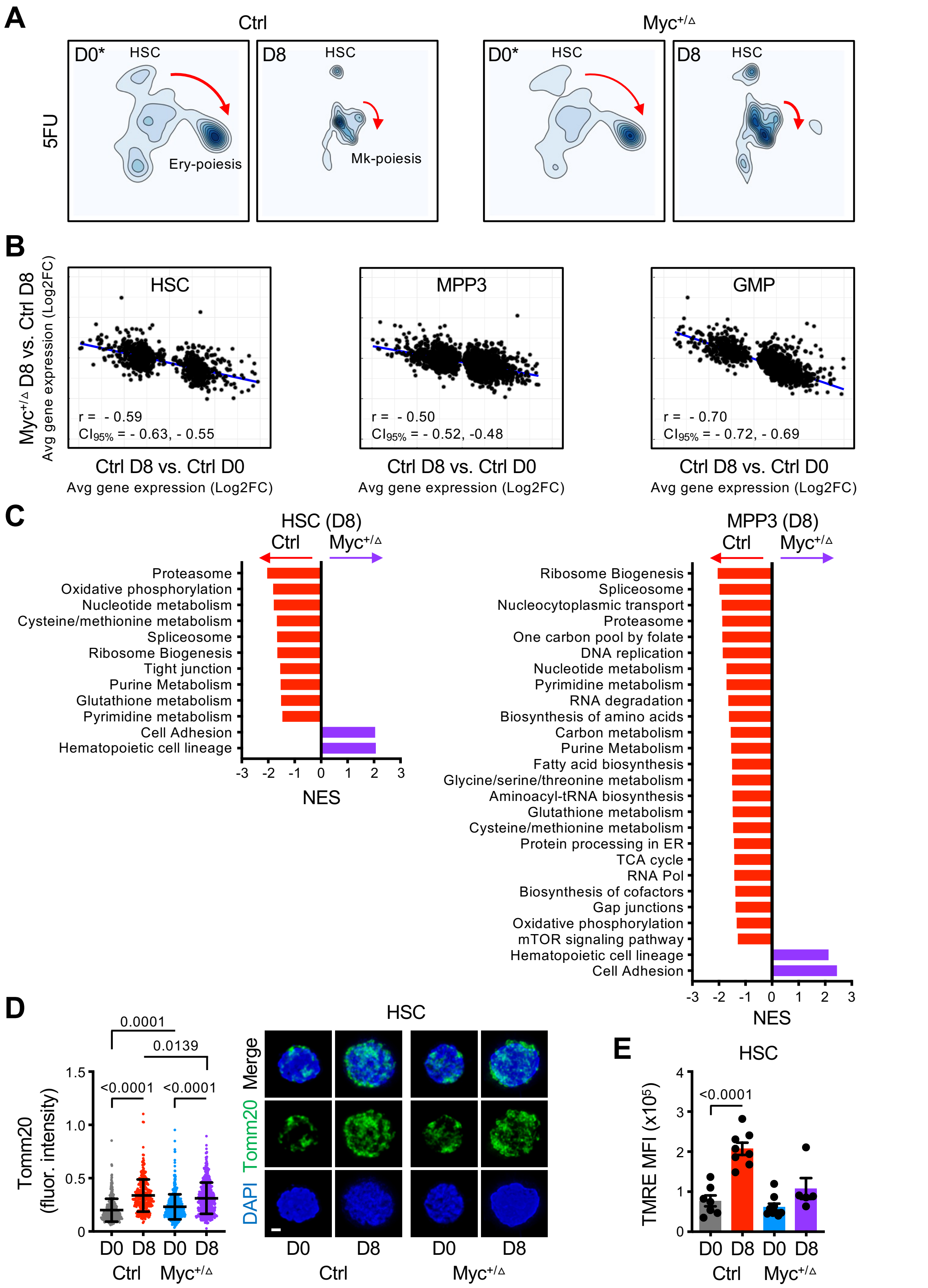


Figure S4 (Olson et al.)

**Figure S4. Impaired transcriptional adaptation in Myc deficient HSPCs, related to Figure 3.**
(A) Transcriptional regulation of regenerative Ctrl and $Myc^{+/\Delta}$ HSPCs post-5FU analyzed by scRNA-seq of BM LK + LSK (D0*) and LK (D8) populations with contour density plots showing the impact of Myc haploinsufficiency on lineage regeneration.
(B) Impact of 5FU treatment on the global transcriptome of Ctrl and $Myc^{+/\Delta}$ HSPCs, with simple linear regression reporting 95% confidence interval of the fitted slope and the R-squared goodness of fit coefficient.
(C) KEGG GSEA analysis of significant ($p$ <0.05) transcriptional changes in Ctrl and $Myc^{+/\Delta}$ HSCs (left) and MPP3s (right) between D0 and D8 post-5FU.
(D) Mitochondria quantification in regenerative Ctrl and $Myc^{+/\Delta}$ HSCs (n=333-511 cells; 3 independent experiments) at D0 and D8 post-5FU analyzed by Tomm20 immunofluorescence, with quantification of Tomm20 fluorescence (fluor.) intensity (left) and representative image (selected from 3 independent experiments; right).
(E) Measurement of mitochondrial membrane potential in regenerative Ctrl and $Myc^{+/\Delta}$ HSCs at D0 and D8 post-5FU by TMRE (n=5-9; 3 independent experiments).
Data are means ± S.D. (D) or ± S.E.M. (E); dots represent individual cells (D) and circles individual mice (E); *P. values* were obtained by the Kolmogorov-Smirnov test (C), and an unpaired t-test (D, E).

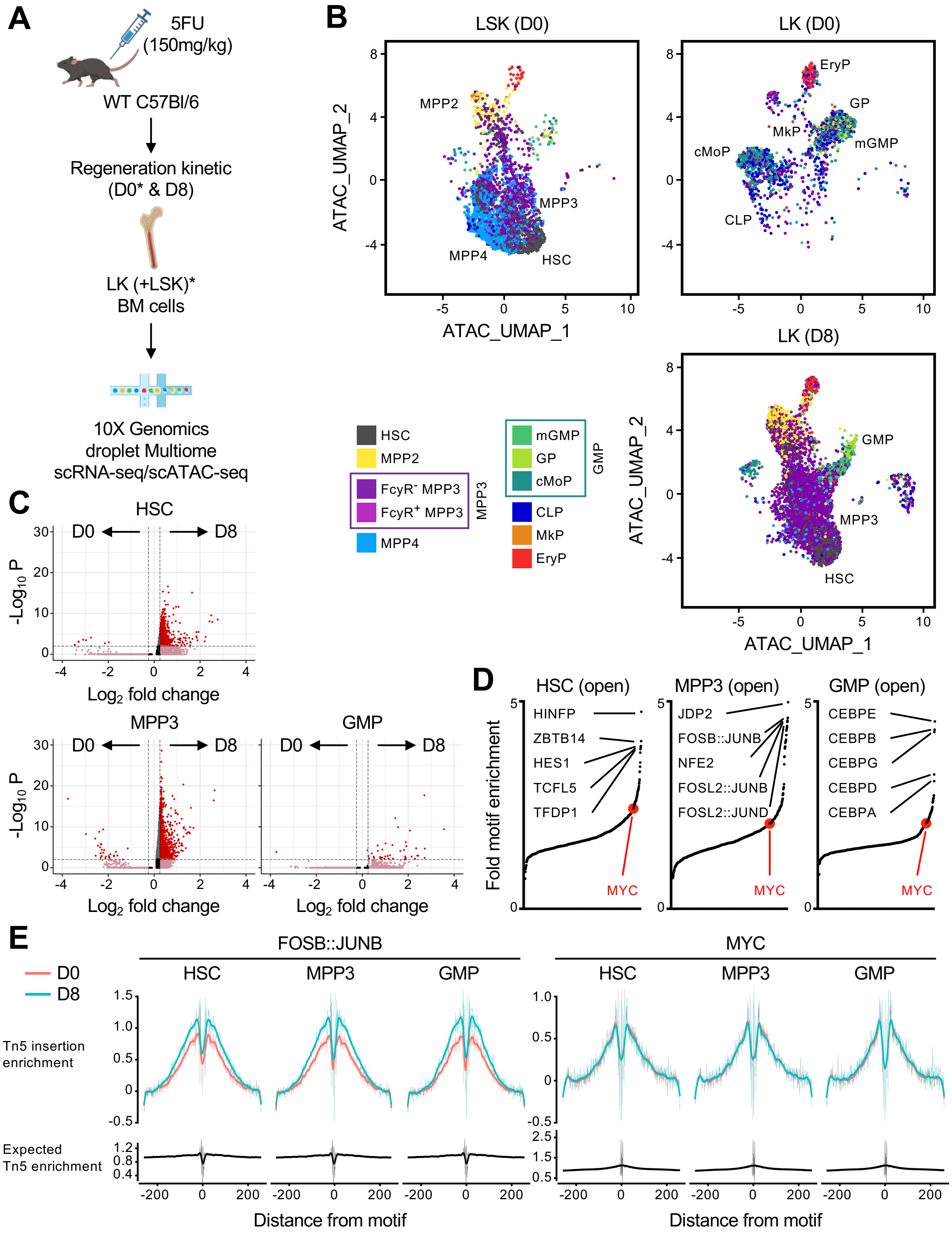


Figure S5 (Olson et al.)

**Figure S5. HSPC chromatin is permissive to Myc hyperactivation, related to Figure 3.**
(A) Chromatin landscape of regenerative HSPCs at D0 and D8 post-5FU by Multiome scRNA-seq/scATAC-seq analysis of BM LSK and LK populations.
(B) UMAP projections of scATAC-seq results with HemaScribe cell type identification generated from the scRNA-seq results.
(C-E) Analyses of D8 vs D0 post-5FU HSPCs: (C) volcano plots of differentially accessible peaks; (D) motif enrichment analysis in opened accessible peaks; and (E) FOSB::JUNB (left) and MYC (right) motif footprints.

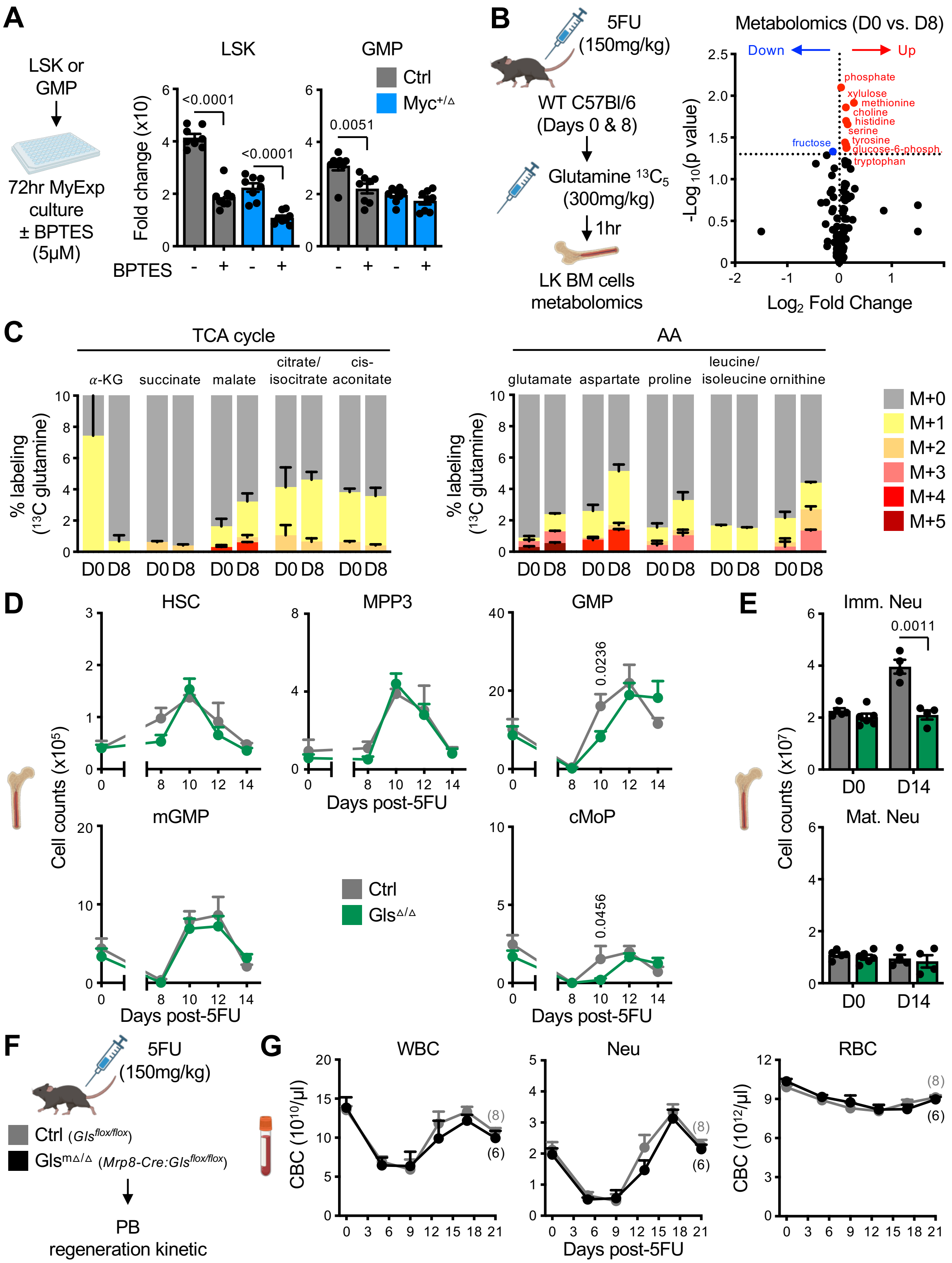


Figure S6 (Olson et al.)

**Figure S6. Glutaminase dependence is specific to EM HSPCs, related to Figure 4.**
(A) Quantification of Ctrl and *Myc*$^{+/\Delta}$ LSK (left) and GMP (right) expansion after 72 hours culture in regenerative-like myeloid expansion conditions with or without (±) the glutamine metabolism inhibitor BPTES (n=8; 4 independent experiments). Results are shown as fold change of plated cells.
(B-C) Metabolism of regenerative HSPCs (n=3; 1 independent experiments): (B) scheme of *in vivo* stable isotope tracing of $C^{13}$-labeled glutamine (1 hour) in BM LK at D0 and D8 post-5FU treatment (left) with volcano plot of LC-MS metabolomic analysis (right); and (C) incorporation of glutamine-derived $C^{13}$ in TCA cycle metabolites (left) and amino acids (AA, right).
(D-E) Regenerative response in hematopoietic-specific glutaminase-deficient mice: (D) quantification of BM HSPC (n=6-16 mice per time point and genotype; 4 independent experiments) post-5FU treatment; and (E) BM $CD101^-$ immature neutrophils (Imm. Neu) and $CD101^+$ mature neutrophils (mat. Neu) at D0 and D14 post-5FU (n=4-6; 2 independent experiments).
(F-G) Regenerative response in myeloid-specific glutaminase-deficient mice: (F) scheme of 5FU treatment in control (Ctrl) and myeloid-specific Gls-deficient mice (*Gls*$^{m\Delta/\Delta}$); and (G) quantification of peripheral blood WBCs, neutrophils, and RBCs (n=6-8; 2 independent experiments).
Data are means ± S.E.M.; circles represent individual mice; *P. values* were obtained by an unpaired t-test (A, D, E).

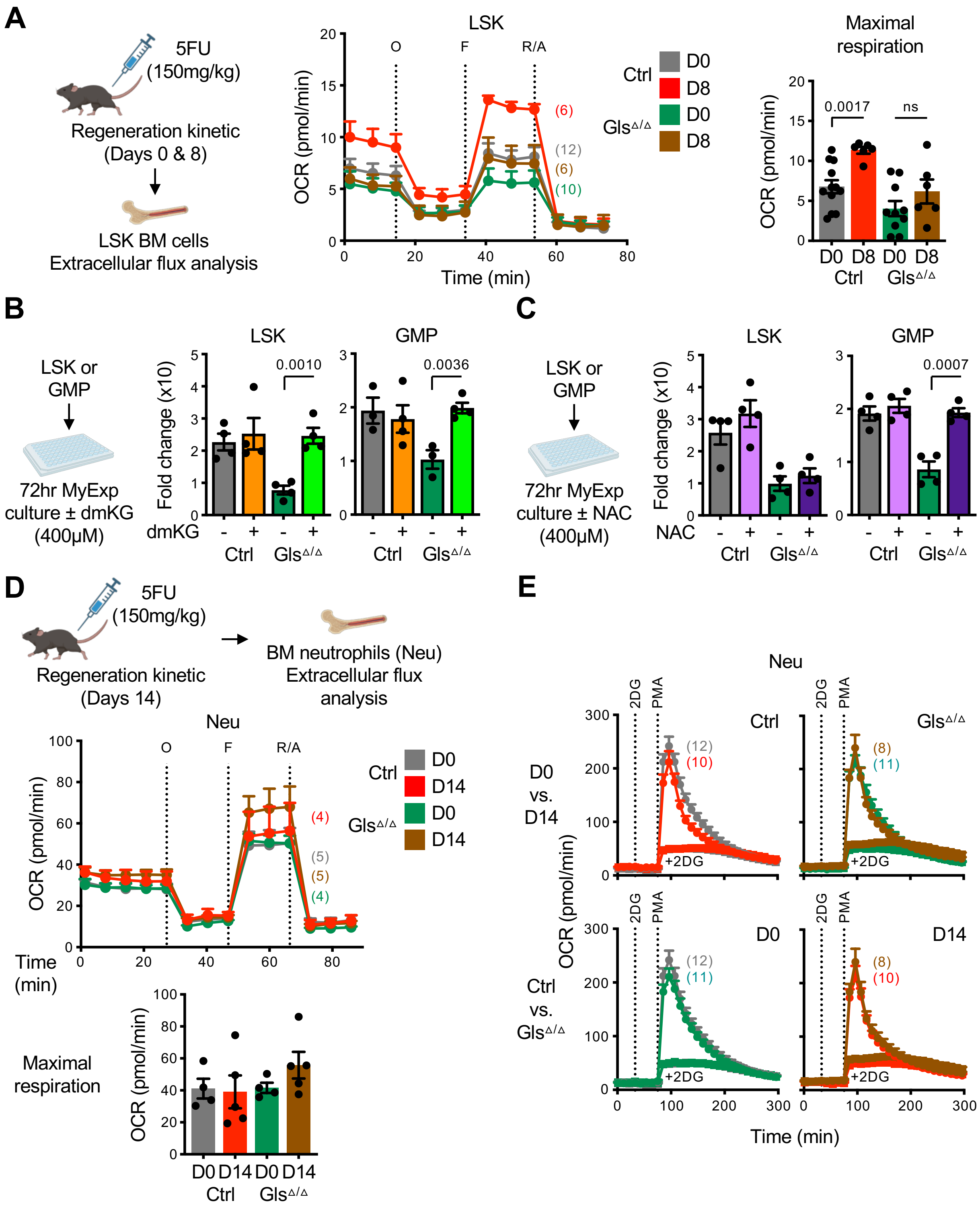


Figure S7 (Olson et al.)

**Figure S7. Anaplerotic glutamine metabolism fuels HSPC OXPHOS during EM, related to Figure 4.**

(A) Scheme of OXPHOS measurement by extracellular flux analysis in regenerative Ctrl and *Gls*$^{\Delta/\Delta}$ LSK at D0 and D8 post-5FU (left), with oxygen consumption rates (OCR) levels (middle) and detailed maximal respiration levels (right) (n=6-12; 6 independent experiments). O, Oligomycin A; F, FCCP; R/A, Rotenone/Antimycin A.

(B-C) In vitro rescue of impaired proliferation from glutaminase-deficient HSPCs: quantification of Ctrl and *Gls*$^{\Delta/\Delta}$ LSK (left) and GMP (right) expansion after 72 hours culture in regenerative-like myeloid expansion conditions with or without (B) the TCA cycle precursor dimethyl alpha-ketoglutarate (dmKG) (n=4; 2 independent experiments) and (C) the antioxidant n-acetylcysteine (NAC) (n=4; 2 independent experiments). Results are shown as fold change of plated cells.

(D) Scheme of OXPHOS measurement by extracellular flux analysis in regenerative Ctrl and *Gls*$^{\Delta/\Delta}$ neutrophils (Neu) at D0 and D14 post-5FU (top), with OCR levels (middle) and detailed maximal respiration levels (bottom) (n=4-5; 4 independent experiments).

(E) Oxidative burst capacity of regenerative Ctrl and *Gls*$^{\Delta/\Delta}$ neutrophils at D0 and D14 post-5FU (n=8-12; 3 independent experiments). PMA, phorbol 12-myristate 13-acetate; 2-DG, 2-Deoxy-d-glucose.

Data are means ± S.E.M.; circles represent individual mice; *P. values* were obtained by an unpaired t-test (A-D).

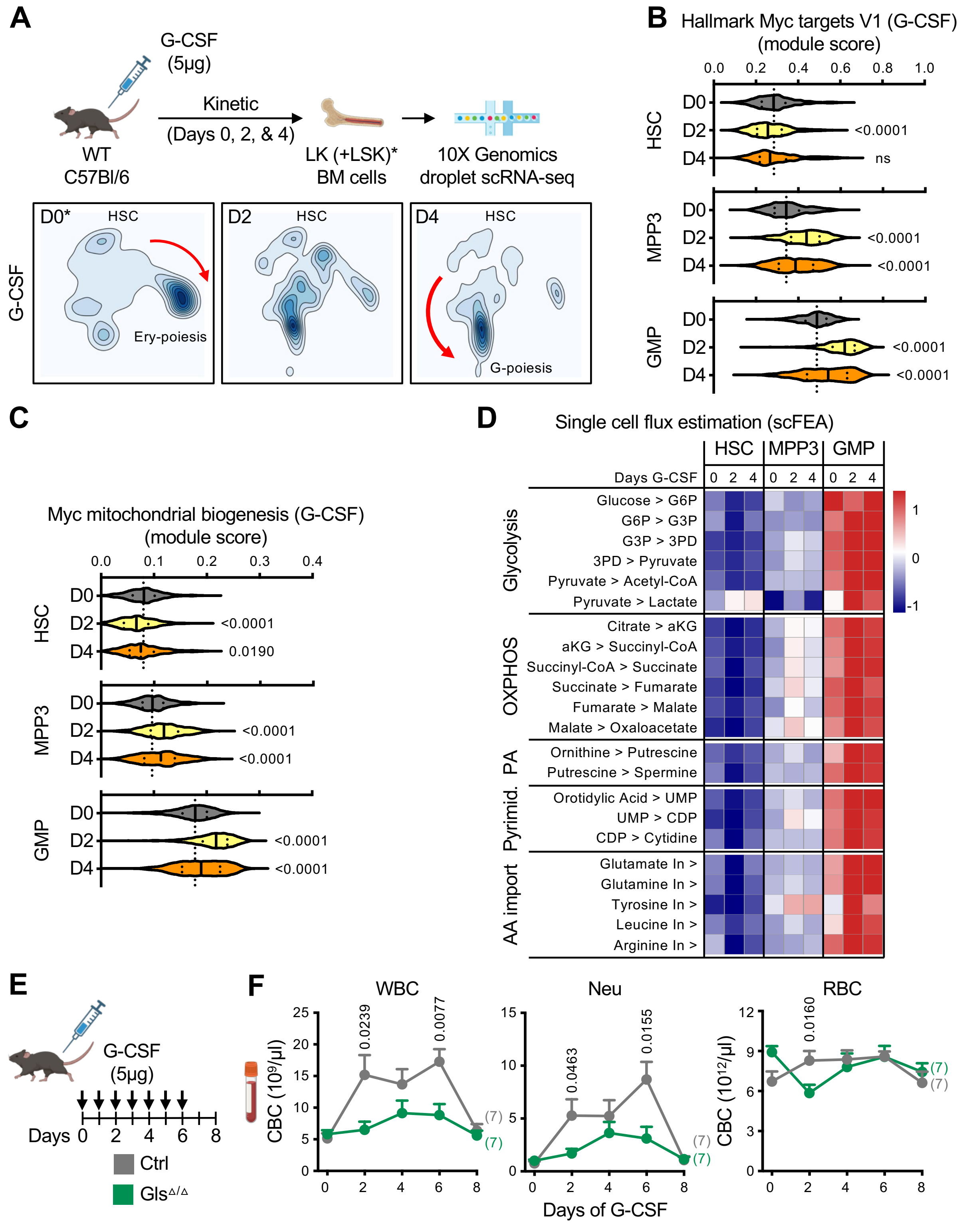


Figure S8 (Olson et al.)

**Figure S8. Myc-driven metabolic adaptation is consistent across EM models, related to Figure 5.**
(A) Transcriptional regulation of regenerative HSPCs following G-CSF treatment analyzed by scRNA-seq of BM LSK and LK populations (top), with contour density plots showing the impact on granulopoiesis.
(B-C) Single-cell assessment of (B) Hallmark Myc V1 transcriptional signature and (C) Myc mitochondrial biogenesis score[27] in HSC, MPP3, and GMP populations during the G-CSF kinetic.
(D) ScFEA analysis of metabolic pathways in G-CSF-treated HSPCs.
(E-F) G-CSF response in hematopoietic-specific glutaminase deficient mice: (E) scheme of G-CSF treatment in control (Ctrl) and *Gls*-deficient ($Gls^{\Delta/\Delta}$) mice; and (F) quantification of peripheral blood WBCs, neutrophils, and RBCs (n=7; 2 independent experiments).
Data are violin plots with quartiles (B, C) or means ± S.E.M. (F); *P. values* were obtained by the Kruskal-Wallis test (B, C) and an unpaired t-test (F).

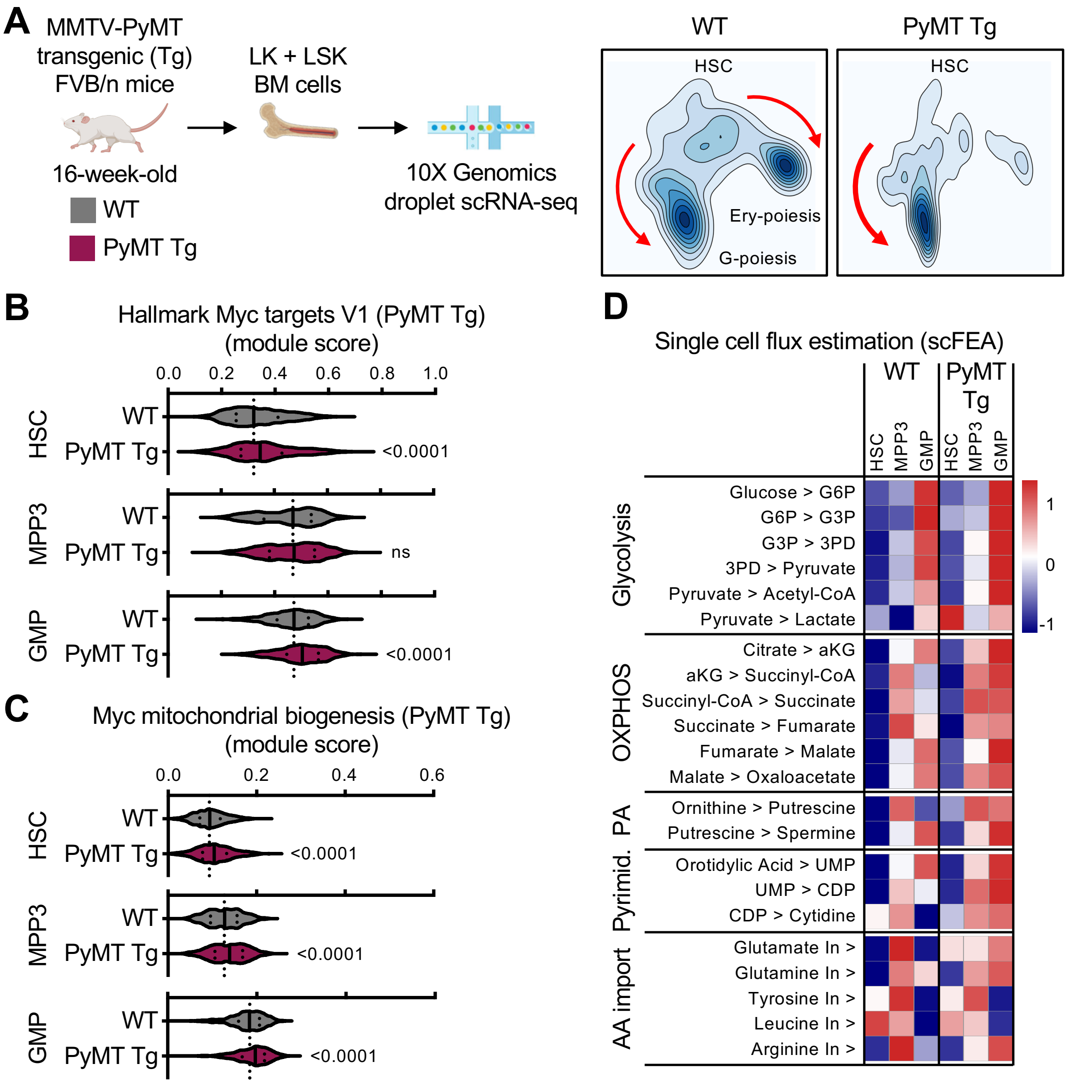


Figure S9 (Olson et al.)

**Figure S9. Myc-driven metabolic adaptation is observed in disease models of breast cancer, related to Figure 5.**
(A) Transcriptional regulation of HSPCs in PyMT transgenic (Tg) model of breast cancer analyzed by scRNA-seq of BM HSPC (LSK + LK) populations (top), with contour density plots showing the impact of tumor burden on granulopoiesis.
(B-C) Single-cell assessment of (B) Hallmark Myc V1 transcriptional signature and (C) Myc mitochondrial biogenesis score[27] in HSC, MPP3, and GMP populations in WT and tumor-bearing PyMT Tg mice.
(D) ScFEA analysis of metabolic pathways in tumor-bearing PyMT Tg HSPCs.
Data are violin plots with quartiles (B, C); *P. values* were obtained by the Kruskal-Wallis test (B, C).

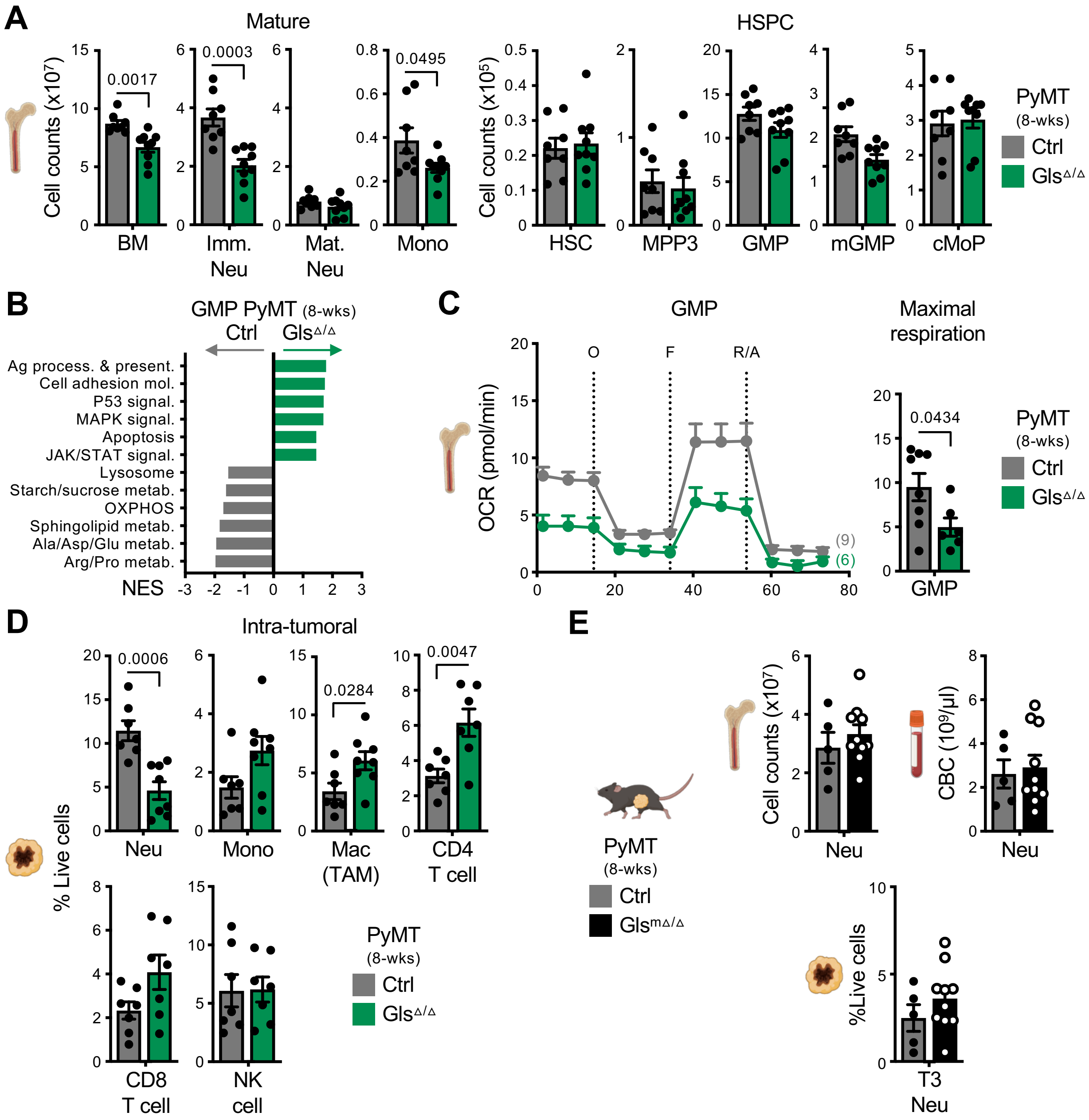


Figure S10 (Olson et al.)

**Figure S10. Hematopoietic glutaminase deficiency alters the composition of the tumor microenvironment, related to Figure 5.**

(A) Quantification of BM mature and HSPC populations 8 weeks after Py230 orthotopic implantation in Ctrl and tumor-bearing $Gls^{\Delta/\Delta}$ mice (n=8-9; 3 independent experiments).

(B) KEGG GSEA analysis of significant ($p$ <0.05) transcriptional changes in GMPs from Ctrl and tumor-bearing $Gls^{\Delta/\Delta}$ mice 8 weeks post-orthotopic implantation.

(C) OXPHOS measurement by extracellular flux analysis of GMP from Ctrl and tumor-bearing $Gls^{\Delta/\Delta}$ mice 8 weeks post-orthotopic implantation, with OCR levels (left) and detailed maximal respiration levels (right) (n=6-9; 3 independent experiments). O, Oligomycin A; F, FCCP; R/A, Rotenone/Antimycin A.

(D) Quantification of intratumoral myeloid and lymphoid populations from Ctrl and tumor-bearing $Gls^{\Delta/\Delta}$ mice 8 weeks post-orthotopic implantation (n=7-8; 3 independent experiments). Neu, neutrophil; Mono, monocyte; Mac, macrophages (TAM); NK, natural killer cells.

(E) Quantification of BM, peripheral blood and intratumoral neutrophil subsets from Ctrl and tumor-bearing myeloid-specific $Gls^{m\Delta/\Delta}$ mice at 8 weeks post-orthotopic implantation (n=6-11; 3 independent experiments).

Data are means ± S.E.M.; circles represent individual mice; *P. values* were obtained by the Kolmogorov-Smirnov test (B), and an unpaired t-test (A, C, D, E).

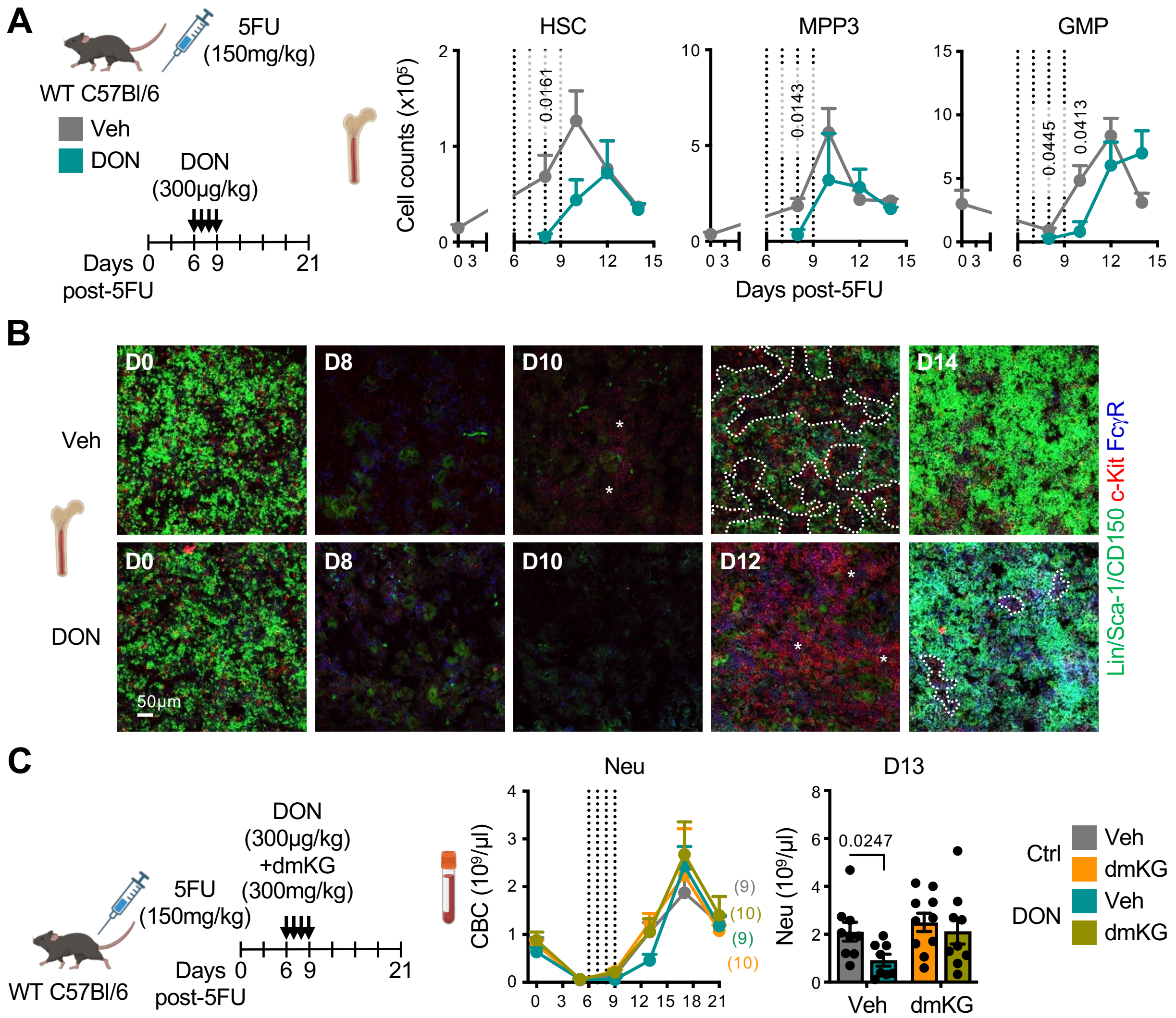


Figure S11 (Olson et al.)

**Figure S11. Pharmacologic inhibition of glutamine metabolism impairs regenerative myelopoiesis, related to Figure 6.**

(A-B) Regenerative response in mice treated with the glutamine metabolism inhibitor 6-Diazo-5-oxo-L-norleucine (DON) or PBS vehicle (Veh) control: (A) BM HSPCs (n=2-7 mice per time point and genotype; 2 independent experiments) post-5FU treatment; and (B) immunofluorescence imaging of BM GMP patches (stars) and GMP clusters (dotted line) post-5FU treatment (representative images from 2 independent experiments).

(C) *In vivo* rescue of impaired regeneration from DON-treated HSPCs: treatment scheme with daily dmKG and DON injections at days 6 to 9 post-5FU treatment in WT mice (n=9-10; 2 independent experiments) (left), quantification of peripheral blood neutrophils (middle), and neutrophil counts at D13 post-5FU (right).

Data are means ± S.E.M.; circles represent individual mice; *P. values* were obtained by an unpaired t-test (A, C).
